# Noncanonical TRAIL Signaling Promotes Myeloid-Derived Suppressor Cell Abundance and Tumor Progression in Cholangiocarcinoma

**DOI:** 10.1101/2023.05.24.541931

**Authors:** Emilien Loeuillard, Binbin Li, Hannah E. Stumpf, Jingchun Yang, Jessica Willhite, Jennifer L. Tomlinson, Juan Wang, Fred Rakhshan Rohakhtar, Vernadette A. Simon, Rondell P. Graham, Rory L. Smoot, Haidong Dong, Sumera I. Ilyas

**Author notes:** Materials & Correspondence: Sumera I. Ilyas, M.B.B.S. Associate Professor of Medicine Mayo Clinic College of Medicine and Science 200 First Street SW Rochester, Minnesota 55905.

## Abstract

Proapoptotic tumor necrosis factor-related apoptosis-inducing ligand (TRAIL) signaling as a cause of cancer cell death is a well-established mechanism. However, TRAIL-receptor (TRAIL-R) agonists have had very limited anticancer activity in humans, challenging the concept of TRAIL as a potent anticancer agent. Herein, we demonstrate that TRAIL^+^ cancer cells can leverage noncanonical TRAIL signaling in myeloid-derived suppressor cells (MDSCs) promoting their abundance in murine cholangiocarcinoma (CCA). In multiple immunocompetent syngeneic, orthotopic murine models of CCA, implantation of TRAIL^+^ murine cancer cells into *Trail-r^-/-^*mice resulted in a significant reduction in tumor volumes compared to wild type mice. Tumor bearing *Trail-r^-/-^* mice had a significant decrease in the abundance of MDSCs due to attenuation of MDSC proliferation. Noncanonical TRAIL signaling with consequent NF-κB activation in MDSCs facilitated enhanced MDSC proliferation. Single cell RNA sequencing and cellular indexing of transcriptomes and epitopes by sequencing (CITE-Seq) of CD45^+^ cells in murine tumors from three distinct immunocompetent CCA models demonstrated a significant enrichment of an NF-κB activation signature in MDSCs. Moreover, MDSCs were resistant to TRAIL-mediated apoptosis due to enhanced expression of cellular FLICE inhibitory protein (cFLIP), an inhibitor of proapoptotic TRAIL signaling. Accordingly, cFLIP knockdown sensitized murine MDSCs to TRAIL-mediated apoptosis. Finally, cancer cell-restricted deletion of *Trail* significantly reduced MDSC abundance and murine tumor burden. In summary, our findings define a noncanonical TRAIL signal in MDSCs and highlight the therapeutic potential of targeting TRAIL^+^ cancer cells for the treatment of a poorly immunogenic cancer.

## INTRODUCTION

Tumor necrosis factor-related apoptosis-inducing ligand (TRAIL), a member of the tumor necrosis factor super family, is predominantly expressed on immune cells, which use it as an important mechanism to induce cancer cell death by apoptosis.^1^ Two TRAIL receptors are expressed in humans: TRAIL-receptor 1 or death receptor 4 (TRAIL-R1/DR4, *TNFRSF10A*) and TRAIL-R2 (DR5, *TNFRSF10B*).^2,3^ Mice only have a single TRAIL receptor (death receptor 5 [Dr5] or Trail-r).^4^ The TRAIL/TRAIL-R system has garnered considerable interest in cancer biology, especially as a potential anticancer therapy.^1^ In multiple murine models of tumorigenesis, *Trail* deficiency renders mice more susceptible to carcinogenesis and tumor progression.^1^ However, TRAIL-R agonists have had very limited anticancer activity in humans,^1^ challenging this concept of TRAIL as an anticancer agent. Indeed, emerging information suggests that TRAIL may have a protumor role as it can foster tumor proliferation as well as invasion.^5-8^ Hence, in addition to having a tumor suppressive role, TRAIL may act as a protumor cytokine that fosters tumor growth and progression, perhaps indirectly by modulating the tumor microenvironment.

The effect of TRAIL/TRAIL-R on the tumor immune microenvironment (TIME) is not well-defined; limited evidence suggests that the influence of TRAIL/TRAIL-R on the TIME is multifaceted and context dependent. The various effects of TRAIL signaling on the TIME include reduction in the abundance of protumor myeloid cells as well as activation of cytotoxic T lymphocytes (CTLs) and natural killer (NK) cells.^1,9-11^ In certain malignancies such as thymoma and breast cancer, TRAIL/TRAIL-R reduces abundance of myeloid-derived suppressor cells (MDSC) by promoting MDSC apoptosis.^10,11^ In contrast, TRAIL induces a promyeloid secretome in lung adenocarcinoma.^5^ Thus, the potency and hierarchy of TRAIL anticancer versus procancer processes in cancer biology has yet to be defined. TRAIL ligation of its cognate receptors induces activation of proapoptotic signaling pathways, and triggers the extrinsic pathway of cellular apoptosis.^12^ Nonapoptotic signaling cascades that promote proinflammatory gene transcription programs and activation of NF-κB have also been described.^13^ However, the biology and pathobiology of noncanonical (nonapoptotic) TRAIL mediated signaling in cancer immunology has not been well defined.

Cholangiocarcinoma (CCA) is a highly lethal desmoplastic cancer originating from the bile ducts.^14^ The majority of CCA patients present with advanced/metastatic disease, and the median overall survival with the current standard-of-care systemic therapy (gemcitabine, cisplatin, and durvalumab) is only 12.8 months.^15^ Immunotherapeutic approaches have had subpar impact on patient survival with low response rates in highly lethal cancers such as CCA. Indeed, the results with immune checkpoint inhibition (ICI) monotherapy in poorly immunogenic tumors such as CCA and pancreatic ductal adenocarcinoma have been disappointing with an overall response rate of <10%,^16,17^ implying that tumor cells ultimately evade the antitumor immune response and further anticancer immune stimulation is necessary. We have demonstrated that the CCA TIME is poorly immunogenic with an abundance of protumor myeloid cells such as tumor-associated macrophages (TAMs) and MDSCs and a paucity of CTLs in the tumor core.^18^ MDSCs are a heterogeneous group of immature myeloid cells that promote immune tolerance, mediate tumor immune escape and immunotherapy resistance with consequent tumor progression.^19^ MDSC subsets include monocytic MDSCs (M-MDSC) that are CD11b^+^Ly6G^−^Ly6C^high^ and granulocytic MDSCs (G-MDSC) that are CD11b^+^Ly6G^+^Ly6C^low^.^19-24^ We have demonstrated that therapeutic targeting of TAMs did not reduce CCA tumor burden in mice due to a compensatory emergence of G-MDSCs, highlighting the integral role of MDSCs in the CCA TIME.^18^

Herein, we examine the role of TRAIL biology in CCA by employing unique immunocompetent murine models. We demonstrate that noncanonical TRAIL signaling promotes an immunosuppressive tumor microenvironment by directly fostering MDSC abundance and function. The protumor, immunosuppressive role of TRAIL/TRAIL-R is potent as implantation of TRAIL^+^ CCA cells into *Trail-r^-/-^* mice or mice with myeloid cell deletion of *Trail-r^-/-^* significantly reduces MDSC abundance and tumor volumes. This noncanonical TRAIL signaling is dependent on the antiapoptotic protein cellular FLICE inhibitory protein (cFLIP) and driven by NF-κB activation in MDSCs. These data suggest that selective antagonism of cancer cell expressed TRAIL can modulate the TIME and reduce tumor immunosuppression.

## RESULTS

### TRAIL-R^+^ immune cells in the TIME are essential in facilitating tumor progression

Whereas TRAIL receptors are ubiquitously expressed, TRAIL is predominantly expressed on cells of the immune system which use it as an important mechanism to induce cancer cell death by apoptosis.^1^ Using single cell RNA sequencing (scRNA seq) of human CCA tumors, we demonstrated that CCA cells also express TRAIL (*TNFSF10*), whereas immune cells express TRAIL-R1 (*TNFRSF10A)* and TRAIL-R2 (*TNFRSF10B*) in human CCA **(Fig. 1A-B)**. Accordingly, the TRAIL/TRAIL-R system may have pleiotropic effects on the CCA TIME. To further examine the role of the TRAIL/TRAIL-R system in the CCA TIME, we leveraged our unique immunocompetent murine models of CCA. These models (SB, KPPC, FAC) employ orthotopic implantation of murine CCA cells and have distinct genetic drivers.^25-27^ Thus, they are an optimal tool for assessing the influence of TRAIL/TRAIL-R on the TIME. Murine CCA cells (SB, KPPC, FAC) have constitutive or cell autonomous TRAIL expression as there is abundant TRAIL protein expression in vitro in these cells despite isolation from the liver **(Supplementary Fig. S1A)**. Murine CCA cells were implanted into livers of WT and *Trail-r^-/-^* C57BL/6J mice. Hence, in this model the host immune cells express TRAIL but not the receptor and would be capable of inducing TRAIL-mediated apoptosis in CCA cells but would be resistant to potential TRAIL-mediated immunosuppression **(Fig. 1C)**. Implantation of SB cells into *Trail-r^-/-^* mice results in a significant reduction in tumor volumes compared to wild type (WT) mice **(Fig. 1C-E)**. Similarly, a significant reduction in tumor volumes was noted in *Trail-r^-/-^* mice compared to WT mice using the KPPC model **(Fig. 1F-H)**. Tumors were also reduced in FAC tumor bearing *Trail-r^-/-^* mice compared to WT mice **(Fig. 1I-K)**. *Trail-r^-/-^* SB, KPPC, and FAC tumors had phenotypic features of CCA **(Supplementary Fig. S1B-D)**. Taken together, these results suggest that TRAIL/TRAIL-R promotes CCA tumor growth, and TRAIL-R^+^ host immune cells are essential in facilitating tumor progression.

**Figure 1.**
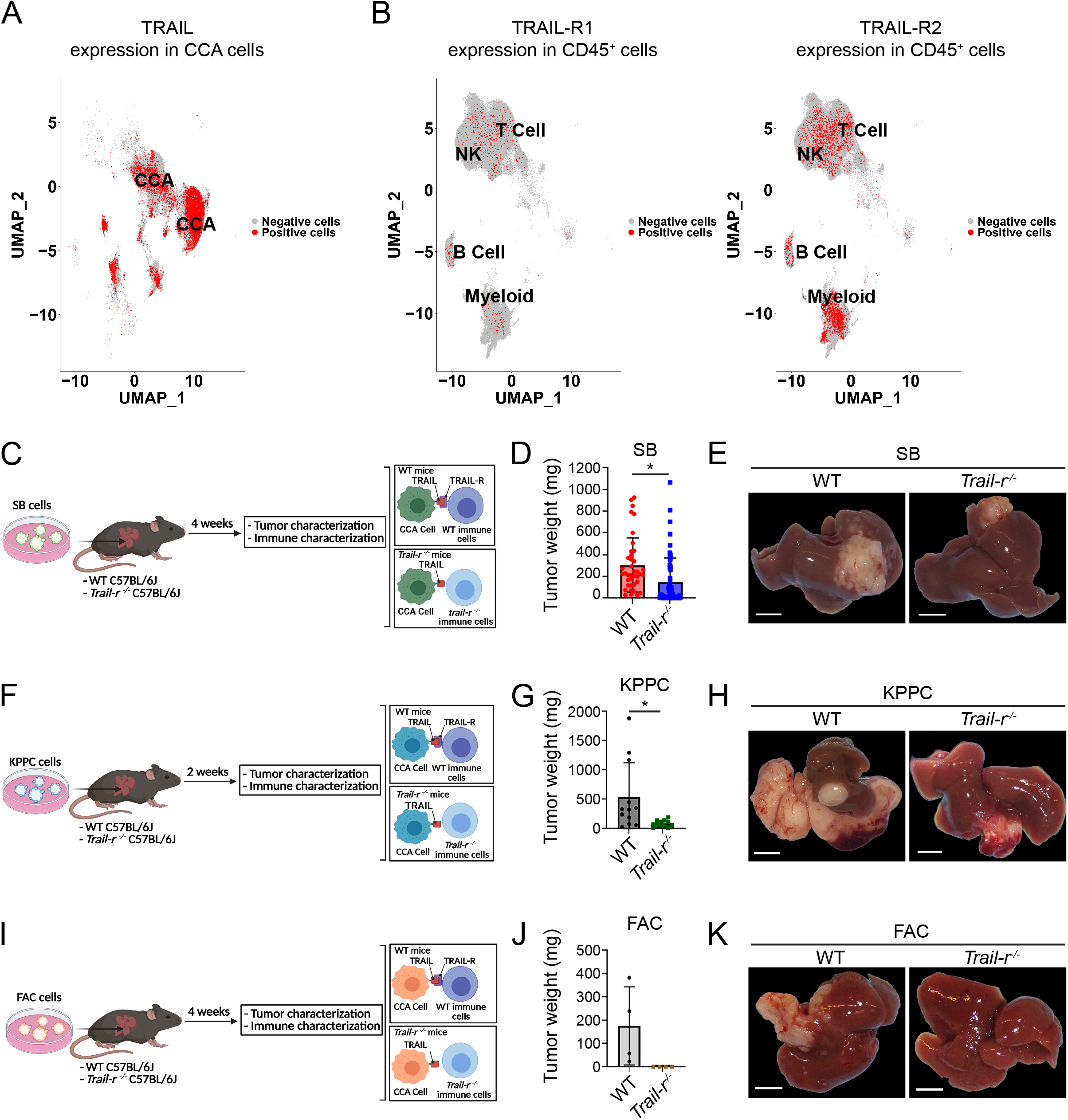
TRAIL-R^+^ immune cells in the TIME are essential in facilitating tumor progression. (**A-B)** UMAP plots of single cells from 34 human CCA tumors in public scRNA seq datasets (GSE210066, GSE189903, GSE138709, GSE125449, HRA000863). **(A)** 7426 CCA cells were colored (red) by TRAIL (*TNFSF10*) expression. **(B)** 156,063 immune cells were colored by TRAIL-R1 (*TNFRSF10A*) expression (left panel) and TRAIL-R2 (*TNFRSF10B*) expression (right panel). Positive cells (red dots) were defined as those with a normalized unique molecular identifier (UMI) of the specific gene larger than 1.7. (**C**) Schematic of SB syngeneic model of CCA in WT and *Trail-r^-l-^* mice. (**D**) Average tumor weight in mg of WT *or Trail-r^-l-^* mice (*n* >10). (**E**) Representative photographs of livers from **D**. Scale bar: 0.5 cm. (**F**) Schematic of KPPC syngeneic model of CCA in WT and *Trail-r^-l-^* mice. (**G**) Average tumor weight in mg of WT *or Trail-r^-l-^* mice (*n* =12). (**H**) Representative photographs of livers from **G**. Scale bar: 0.5 cm. (**I**) Schematic of FAC syngeneic model of CCA in WT and *Trail-r^-l-^* mice. (**J**) Average tumor weight in mg of WT *or Trail-r^-l-^* mice (*n* = 4). (**K**) Representative photographs of livers from **J**. Scale bar: 0.5 cm. Data are represented as mean ± SD. Unpaired Student’s *t* test was used. \**P* < 0.05.

### Genetic deletion of *Trail-r* augments CTL infiltration and function

Tumors arising in mice genetically deficient in *Trail-r* exhibited enhanced CD8^+^ T cell infiltration compared to WT SB and KPPC mice tumors **(Fig. 2A-B; Supplementary Fig. S2A)**. Reactive CTLs express CD11a, an integrin, which is upregulated in effector and memory CD8^+^ T cells. CD11a mediates conjugation between CTLs and target cells and can be used to identify and monitor endogenous tumor-reactive CTLs.^28,29^ *Trail-r^-/-^*tumors had a significant increase in tumor-reactive CD8^+^ T cells and had enhanced CTL effector function **(Fig. 2C-E)**. There was no significant difference in the abundance of CD4^+^ T cells in *Trail-r^-/-^* tumors compared to WT tumors, albeit there was a significant reduction in NK cells in *Trail-r^-/-^* tumors **(Supplementary Fig. S2B, S2C)**. The TRAIL/TRAIL system has been implicated in T cell homeostasis.^30^ Thus, we postulated that TRAIL/TRAIL-R may directly impact CTL apoptosis or proliferation. Although we did not observe a decrease in CTL apoptosis, an increase in proliferation was noted in *Trail-r^-/-^* tumors **(Fig. 2F, 2G)**. To further examine this, murine T cells were isolated from WT or *Trail-r^-/-^*mice and cocultured with CCA (SB) cells **(Fig. 2H)**. No difference in T cell proliferation or apoptosis was noted between WT CD8^+^ T cells and *Trail-r^-/-^* CD8^+^ T cells when cocultured with murine CCA cells, indicating that TRAIL/TRAIL-R does not have a direct effect on T cell proliferation or apoptosis **(Fig. 2I)**. Moreover, in the presence of murine SB cells, WT CD8^+^ T cells had no change in markers of activation, effector function, or exhaustion when compared to *Trail-r^-/-^* CD8^+^ T cells **(Supplementary Fig. S2D-2G)**. Finally, to definitively determine whether TRAIL/TRAIL-R has a direct impact on CD8^+^ T cells and CCA tumor growth, we generated mice with CD8^+^ T cell specific knockout of the *Trail-r* gene by crossing a mouse floxed for the *Trail-r* gene with a Cd8a-cre mouse, a transgenic mouse with Cre activity noted in peripheral CD8^+^ T cells **(Supplementary Fig. S2H)**.^31^ SB cells were implanted orthotopically into livers of *Trail-r^f/f^* and Cd8^cre*-*^*Trail-r^f/f^* mice (ΔCd8) **(Fig. 2J)**. There was no change in SB tumor weights in ΔCd8 mice compared with *Trail-r^f/f^* mice **(Fig. 2K and L)**. Tumors arising in ΔCd8 mice had phenotypic features of CCA **(Supplementary Fig. S2I)**. These data suggest that TRAIL/TRAIL-R does not have a direct effect on CD8^+^ T cells, and the increased CTL abundance in *Trail-r^-/-^* mice may be an indirect effect in the CCA TIME via modulation of immunosuppressive cells by TRAIL/TRAIL-R.

**Figure 2.**
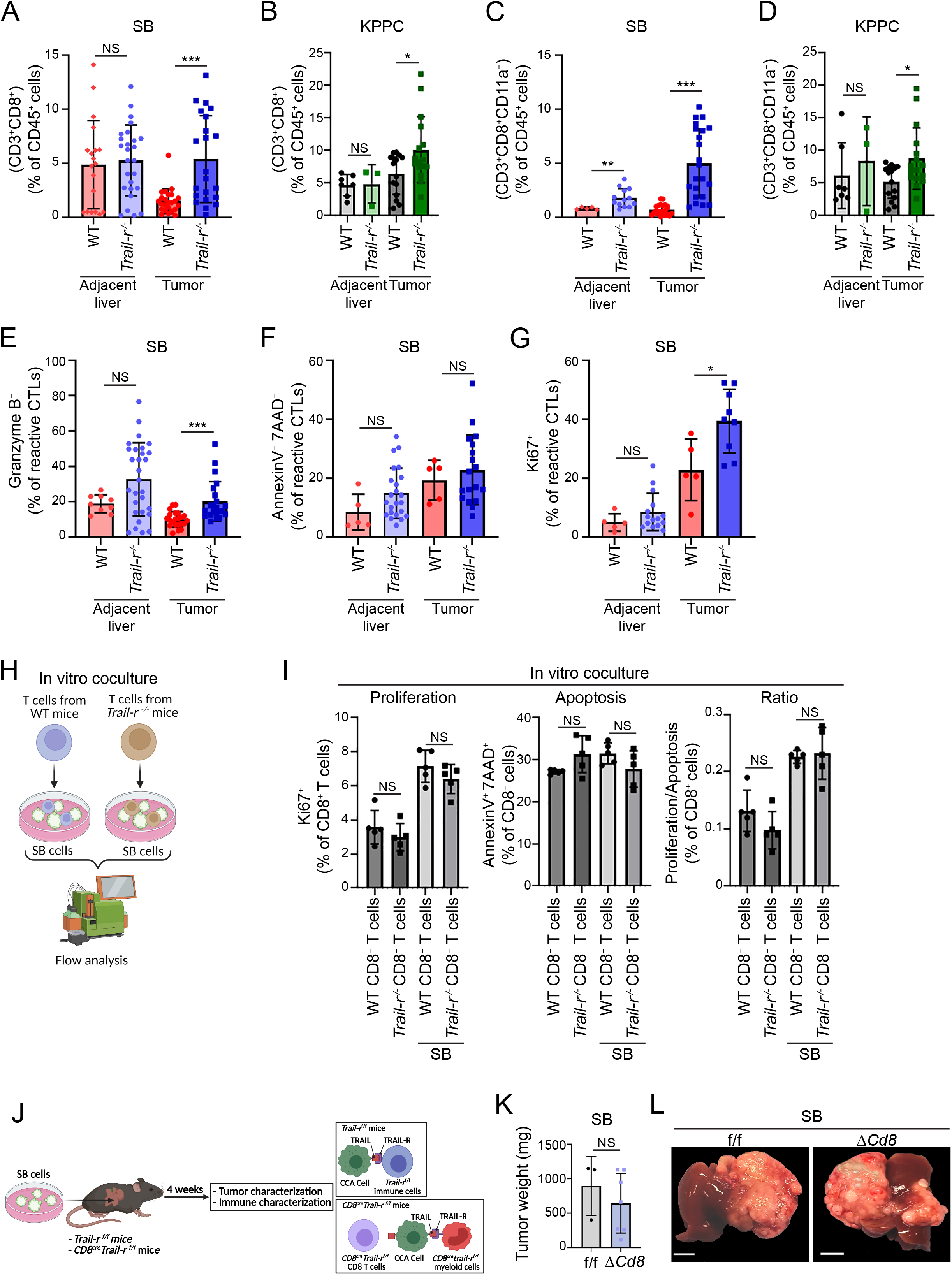
Genetic deletion of *Trail-r* augments CTL infiltration and function. (**A**–**G**) Immune profiling of tumors and tumor-adjacent livers 28 days after orthotopic implantation of 0.75×10^6^ SB cells or 14 days after orthotopic implantation of 0.75×10^6^ KPPC cells in WT or *Trail-r^-/-^* mouse livers. (**A**) Percentage of CTLs (CD45^+^CD3^+^CD8^+^) of CD45^+^ cells in tumor-adjacent livers or SB tumors from WT and *Trail-r^-/-^* mice (*n* ≥ 5). (**B**) Percentage of CTLs (CD45^+^CD3^+^CD8^+^) of CD45^+^ cells in tumor-adjacent livers or KPPC tumor from WT and *Trail-r^-/-^* mice (*n* ≥ 3). (**C**) Percentage of reactive CTLs (CD45^+^CD3^+^CD8^+^CD11a^+^) of CD45^+^ cells in tumor-adjacent livers or SB tumors from WT and *Trail-r^-/-^* mice (*n* ≥ 5). (**D**) Percentage of reactive CTLs (CD45^+^CD3^+^CD8^+^CD11a^+^) of CD45^+^ cells in tumor-adjacent livers or KPPC tumors from WT and *Trail-r^-/-^* mice (*n* ≥ 3). (**E**) Percentage of granzyme B expressed in CD8^+^CD11a^+^ reactive CTLs (CD45^+^CD3^+^CD8^+^CD11a^+^) in tumor-adjacent livers or SB tumors from WT and *Trail-r^-/-^* mice (*n* ≥ 9). (**F**) Percentage of Annexin V^+^7AAD^+^ CTLs in tumor-adjacent livers or SB tumors from WT and *Trail-r^-/-^* mice (*n* ≥ 5). (**G**) Percentage of Ki67^+^ CTLs in tumor-adjacent livers or SB tumors from WT and *Trail-r^-/-^* mice (*n* ≥ 5). (**H**) Schematic of T cell and SB cell coculture experiment. Cells were cocultured for 36 hours. (**I**) Percentage of Ki67^+^ CD8^+^ T cells (CD3^+^CD8^+^) (right panel), percentage of Annexin V^+^7AAD^+^ CD8^+^ T cells (CD3^+^CD8^+^) (middle panel), and ratio of proliferation/apoptosis (left panel) *n* ≥ 5). (**J**) Schematic of CD8cre-Trail*-r^f/f^* (ΔCD8) model of CCA implanted with SB cells. (**K**) Average tumor weight in mg of f/f *or* ΔCD8 mice (*n* ≥ 3). (**L**) Representative photographs of livers from **K**. Scale bar: 0.5 cm. Data are represented as mean ± SD. Unpaired Student’s *t* test was used. ns=nonsignificant; \**P* < 0.05; \*\*\**P* < 0.001.

### Genetic deletion of *Trail-r* restricts murine tumor progression via a significant reduction in MDSCs

As TRAIL signaling did not appear to have a direct suppressive effect on cytotoxic elements within the CCA TIME, we next examined the immunosuppressive cell types. *Trail-r^-/-^* tumors exhibited a significant reduction in MDSCs and macrophages **(Fig. 3A; Supplementary Fig. S3A-B)**, while there was no difference in the abundance of dendritic cells (**Supplementary Fig. S3C)**. *Trail-r^-/-^* SB tumors had a significant attenuation of CD11b^+^Ly6G^+^Ly6C^-^ G-MDSCs without a significant alteration of CD11b^+^Ly6G^-^ Ly6C^+^ M-MDSCs **(Fig. 3A)**. *Trail-r^-/-^* SB tumors also had a significant decrease in CD11b^+^F4/80^+^ macrophages compared to WT SB tumors. CD206^+^ TAMs were decreased in *Trail-r^-/-^* SB tumors, whereas M1-like macrophages, which are typically antitumor and proinflammatory, were significantly increased **(Supplementary Fig. S3A and S3B)**. These data support the notion that *Trail-r^-/-^*SB tumors have a proinflammatory, antitumor milieu which contributes to a significant reduction in tumor volumes in the context of genetic *Trail-r* deficiency. In comparison, *Trail-r^-/-^* KPPC tumors had a significant decrease in M-MDSCs rather than G-MDSCs **(Fig. 3B).** Additionally, *Trail-r^-/-^* KPPC tumors had no significant change in macrophages (**Supplementary Fig. S3D and S3E)**, which is consistent with our prior observation that SB and KPPC tumors have distinct immune microenvironments, yet both models demonstrated a significant decrease in MDSCs in the context of *Trail* deficiency.

**Figure 3.**
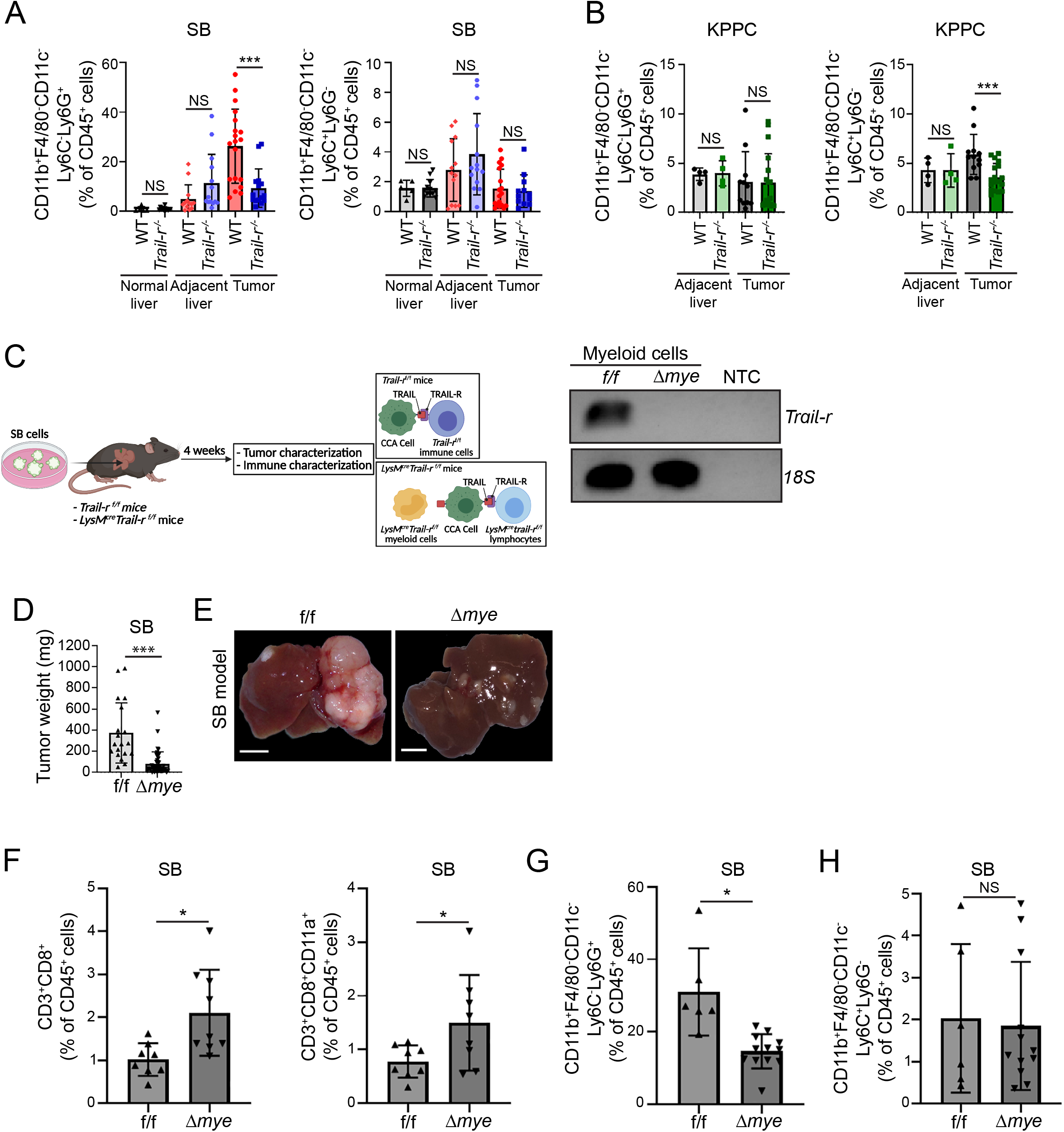
Genetic deletion of *Trail-r* restricts murine tumor progression via a significant reduction in MDSCs. (**A**) Percentage of G-MDSCs (CD45^+^F4/80^-^CD11b^+^CD11c^-^Ly6C^-^Ly6G^+^) of CD45^+^ cells (left panel) and M-MDSCs (CD45^+^F4/80^-^CD11b^+^CD11c^-^Ly6C^+^Ly6G^-^) of CD45^+^ cells (right panel) in WT mouse livers from mice without tumors, tumor-adjacent livers and SB tumors from WT and *Trail-r^-/-^* mice (*n* ≥ 5). (**B**) Percentage of G-MDSCs (left panel) and M-MDSCs (right panel) in tumor-adjacent livers and KPPC tumors from WT and *Trail-r^-/-^* mice (*n* ≥ 4). (**C**) Schematic of *LysM^cre^Trail-r^f/f^* mouse model of CCA and *Trail-r^f/f^* mice. Expression of *Trail-r* in MDSCs differentiated from bone marrow of *Trail-r^f/f^*and *LysM^cre^Trail-r^f/f^* (Δmye) mice. RT-PCR was used to validate *Trail-r* deletion in mice using a primer pair targeting the deletion region in cDNA after Cre-flox editing. 18S rRNA was used as a normalization control. Water was used as a no template control (NTC). (**D**) Average tumor weight in mg of f/f or Δmye mice (*n* ≥ 8). (**E**) Representative photographs of livers from **D**. Scale bar: 0.5 cm. (**F**) Percentage of CTLs (CD45^+^CD3^+^CD8^+^) and reactive CTLs (CD45^+^CD3^+^CD8^+^CD11a^+^) of CD45^+^ cells in SB tumors from f/f and Δmye mice (*n* ≥ 8). (**G**) Percentage of G-MDSCs of CD45^+^ cells in SB tumors from f/f and Δmye mice (*n* ≥ 6). (**H**) Percentage of M-MDSCs of CD45^+^ cells in SB tumors from f/f and Δmye mice (*n* ≥ 6). Data are represented as mean ± SD. Unpaired Student’s *t* test was used. ns=nonsignificant; \**P* < 0.05; \*\*\**P* < 0.001.

As we have previously demonstrated that MDSCs and TAMs play an integral role in the progression of CCA,^18^ we postulated that TRAIL/TRAIL-R promotes tumor progression by mediating tumor immune evasion via these immunosuppressive myeloid cells. To further examine this concept, we employed myeloid cell restricted deletion of *Trail-r*. We generated mice with myeloid cell specific knockout of the *Trail-r* gene by crossing a mouse floxed for the *Trail-r* gene with a *LysM^cre^* mouse.^32^ SB cells were implanted orthotopically into livers of *Trail-r ^f/f^* (f/f) and *LysM^cre^-Trail-r^f/f^* mice (Δmye) **(Fig. 3C)**. There was a significant reduction in SB tumor weights in Δmye mice compared with f/f mice **(Fig. 3D and E)**. Histopathological assessment confirmed that Δmye SB tumors had phenotypic features of CCA **(Supplementary Fig. S3F)**. Similar to our observations in *Trail-r^-/-^* SB tumors, Δmye SB tumors had a significant increase in CD8^+^ T cells and tumor-reactive CD8^+^ T cells compared to f/f SB tumors **(Fig. 3F)**. Consistently, G-MDSCs were also significantly reduced in Δmye SB tumors while no change was observed in M-MDSCs **(Fig. 3G and H)**. In contrast to *Trail-r^-/-^* SB tumors, myeloid cell restricted deletion of *Trail-r* did not influence macrophage abundance, suggesting that MDSCs are the cell type primarily impacted by TRAIL/TRAIL-R (**Supplementary Fig. S3G and S3H)**. In aggregate, these observations confirm that TRAIL/TRAIL-R promotes tumor progression by decreasing tumor infiltrating CTLs, likely via augmentation of MDSC abundance.

### TRAIL/TRAIL-R augments MDSCs abundance via enhanced cellular proliferation

Next, we sought to clarify whether TRAIL/TRAIL-R has a direct effect on MDSC abundance via enhanced proliferation or attenuation of apoptosis. G-MDSCs in *Trail-r^-/-^* tumors had no change in apoptosis, suggesting that these cells are resistant to TRAIL-mediated apoptosis **(Fig. 4A)**. G-MDSCs in *Trail-r^-/-^* tumors did exhibit a significant suppression of proliferation, as assessed by Ki67, compared to MDSCs from WT SB tumors **(Fig. 4B)**. These findings were confirmed in tumor bearing Δmye mice; G-MDSCs had a significant suppression of proliferation without a change in apoptosis in the context of myeloid restricted deletion of *Trail-r* **(Fig. 4C-D)**. M-MDSC abundance was not altered in *Trail-r^-/-^* SB tumors **(Fig. 3A)**. Accordingly, no change was noted in apoptosis or proliferation of M-MDSCs in *Trail-r^-/-^* SB tumors or Δmye SB tumors **(Supplementary Fig. S4A-B)**. Additionally, the percentage of arginase 1 or IL-10 expressing G-MDSCs or M-MDSCs was similar between WT, *Trail-r^-/-^*, and Δmye tumors **(Supplementary Fig. S4C-S4F)**. Overall, these results suggest that TRAIL signaling augments G-MDSC abundance in SB tumors via enhanced proliferation, rather than augmenting expression of immunosuppressive mediators.

**Figure 4.**
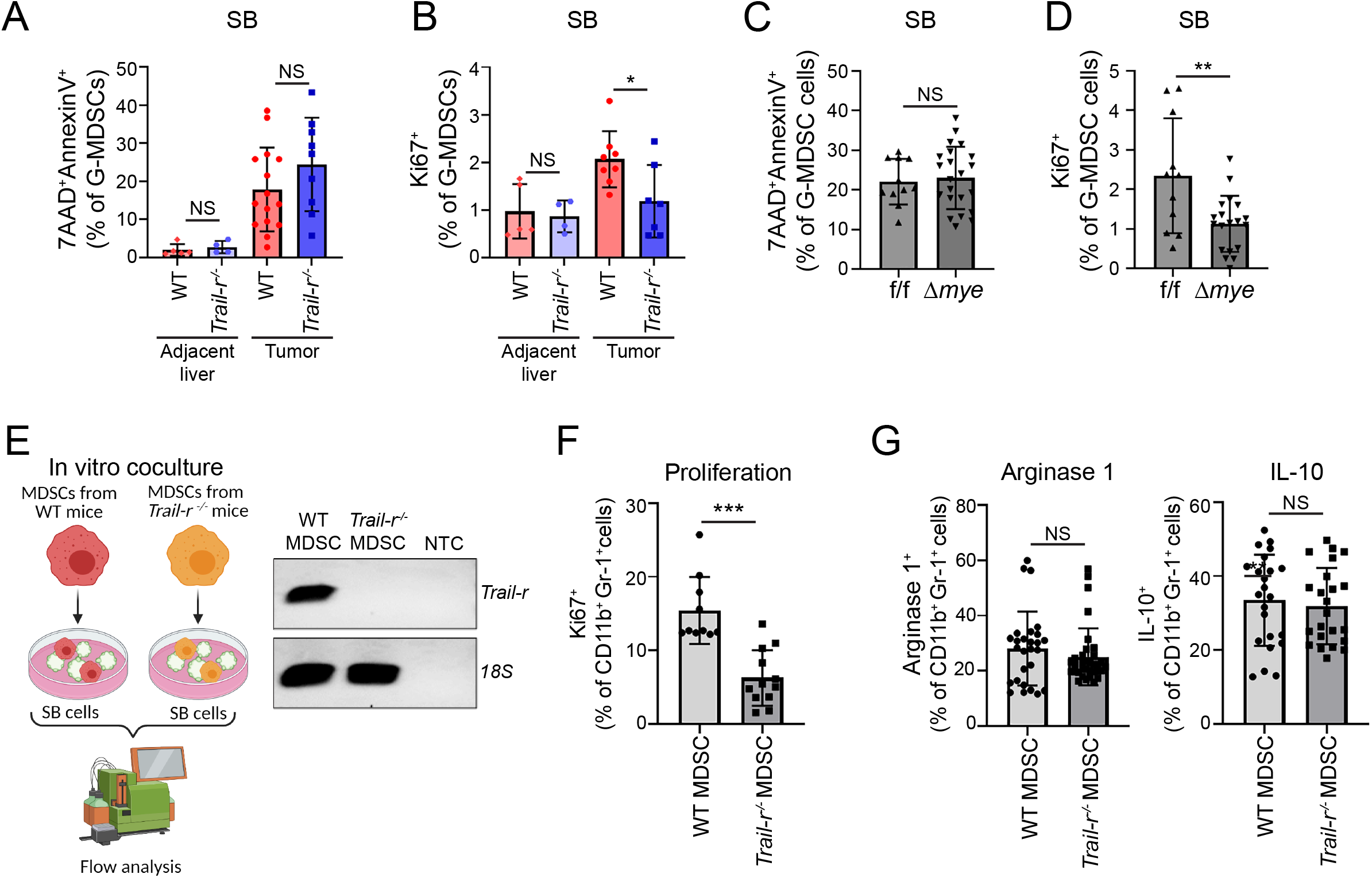
TRAIL/TRAIL-R augments MDSCs abundance via enhanced proliferation. (**A**) Percentage of Annexin V^+^7AAD^+^ G-MDSCs (CD45^+^F4/80^-^CD11b^+^CD11c^-^Ly6C^-^Ly6G^+^) of CD45^+^ cells in tumor-adjacent livers and SB tumors from WT and *Trail-r^-/-^* mice (*n* ≥ 5). (**B**) Percentage of Ki67^+^ G-MDSCs (CD45^+^F4/80^-^CD11b^+^CD11c^-^Ly6C^-^Ly6G^+^) of CD45^+^ cells in tumor-adjacent livers and SB tumors from WT and *Trail-r^-/-^* mice (*n* ≥ 4). (**C**) Percentage of Annexin V^+^7AAD^+^ G-MDSCs (CD45^+^F4/80^-^ CD11b^+^CD11c^-^Ly6C^-^Ly6G^+^) of CD45^+^ cells in tumor-adjacent livers and SB tumors from f/f and Δmye mice (*n* ≥ 11). (**D**) Percentage of Ki67^+^ G-MDSCs (CD45^+^F4/80^-^CD11b^+^CD11c^-^Ly6C^-^Ly6G^+^) of CD45^+^ cells in tumor-adjacent livers and SB tumors from f/f and Δmye mice (*n* ≥ 11). (**E**) Schematic of in vitro coculture of SB cells and BM-derived MDSCs from WT and *Trail-r* mice. Expression of *Trail-r* in MDSCs differentiated from bone marrow of WT and *Trail-r* mice. 18s rRNA was used as a normalization control. Water was used as a no template control (NTC). (**F**) Percentage of Ki67^+^ MDSCs (CD11b^+^Gr-1^+^) after 24h of coculture with SB cells (*n* ≥ 10). (**G**) Percentage of Arginase 1^+^ MDSCs (CD11b^+^Gr-1^+^) (left panel) and IL-10^+^ MDSCs (right panel) after 24h of coculture with SB cells (*n* ≥ 10). Data are represented as mean ± SD. Unpaired Student’s *t* test was used. ns=nonsignificant; \**P* < 0.05; \*\**P* < 0.01; \*\*\**P* < 0.001.

To determine whether enhanced MDSC proliferation is a direct effect of TRAIL signaling activation, we differentiated CD11b^+^Ly6C^+^ MDSCs from bone marrow (BM) of WT and *Trail-r^-/-^* mice using growth factors as previously described.^33^ We confirmed expression of *Trail-r* in WT BM-derived MDSCs (WT MDSCs) and knockout of *Trail-r* in MDSCs from *Trail-r^-/-^* mice (*Trail-r*^-/-^ MDSCs) **(Fig. 4E)**. WT and *Trail-r^-/-^* MDSCs were cocultured with murine tumor cells (SB cells) that express TRAIL protein (**Supplementary Fig. S1A)** and MDSCs were subsequently harvested for further analysis via flow cytometry **(Fig. 4E)**. *Trail-r*^-/-^ MDSCs had a significant decrease in proliferation compared to WT MDSCs when cocultured with tumor cells **(Fig. 4F)**, suggesting that TRAIL^+^ cancer cells bind to TRAIL-R on MDSCs and directly influence MDSC abundance by increasing MDSC proliferation. Moreover, no change was observed in the expression of markers of immunosuppression on WT MDSCs or *Trail-r*^-/-^ MDSCs when cocultured with tumor cells **(Fig. 4G)**, suggesting that the primary effect is on MDSC abundance rather than function. Collectively, these data demonstrate that G-MDSCs are resistant to TRAIL-mediated apoptosis in murine CCA tumors and TRAIL/TRAIL-R directly influences MDSC abundance by increasing MDSC proliferation.

### Noncanonical TRAIL-R signaling promotes MDSC survival in an NF-κB dependent manner

Activation of the classical NF-κB pathway leads to phosphorylation and degradation of inhibitor of κB (IκB) kinase complex which liberates cytosolic NF-κB with subsequent nuclear translocation of NF-κB subunits p50 and p65.^34^ We have observed that TRAIL signaling augments MDSC proliferation **(Fig. 4B)**. To delineate whether this observed effect of TRAIL signaling on MDSC proliferation is occurring via noncanonical TRAIL signaling, we examined NF-κB activation in MDSCs. Addition of SB conditioned medium to BM-derived MDSCs resulted in a significant, time-dependent increase in IκBα phosphorylation and nuclear translocation of the p65 subunit of NF-κB (**Supplementary Fig. S5A-B).** Notably, *Trail-r^-/-^* MDSCs had a significant decrease in IκBα phosphorylation and in nuclear translocation of p65 compared to WT MDSCs when incubated with tumor cell conditioned medium, suggesting that TRAIL/TRAIL-R may augment MDSC abundance via NF-κB activation (**Fig. 5A-B**). Next, we examined the impact of NF-κB signaling inhibition using a small molecule inhibitor of IKKα/β, TPCA-1,^35^ on murine MDSC proliferation. TPCA-1 significantly reduced proliferation of WT MDSCs cocultured with SB cells (**Fig. 5C-D**). TPCA-1 also reduced nuclear translocation of NF-κB of WT MDSCs cocultured with SB cells (data not shown). We then conducted scRNA seq and cellular indexing of transcriptomes and epitopes by sequencing (CITE-Seq) of CD45^+^ cells isolated from murine SB, KPPC, and FAC tumors. Unsupervised clustering revealed 11 clusters of immune cells in the murine tumors (**Supplementary Fig. S5C**). Significant enrichment of a previously reported NF-κB activation signature^36^ was noted in the G-MDSC cluster (**Fig. 5E**). Collectively, these data indicate that noncanonical TRAIL signaling will result in increased NF-κB activation in MDSCs with consequent enhanced proliferation.

**Figure 5.**
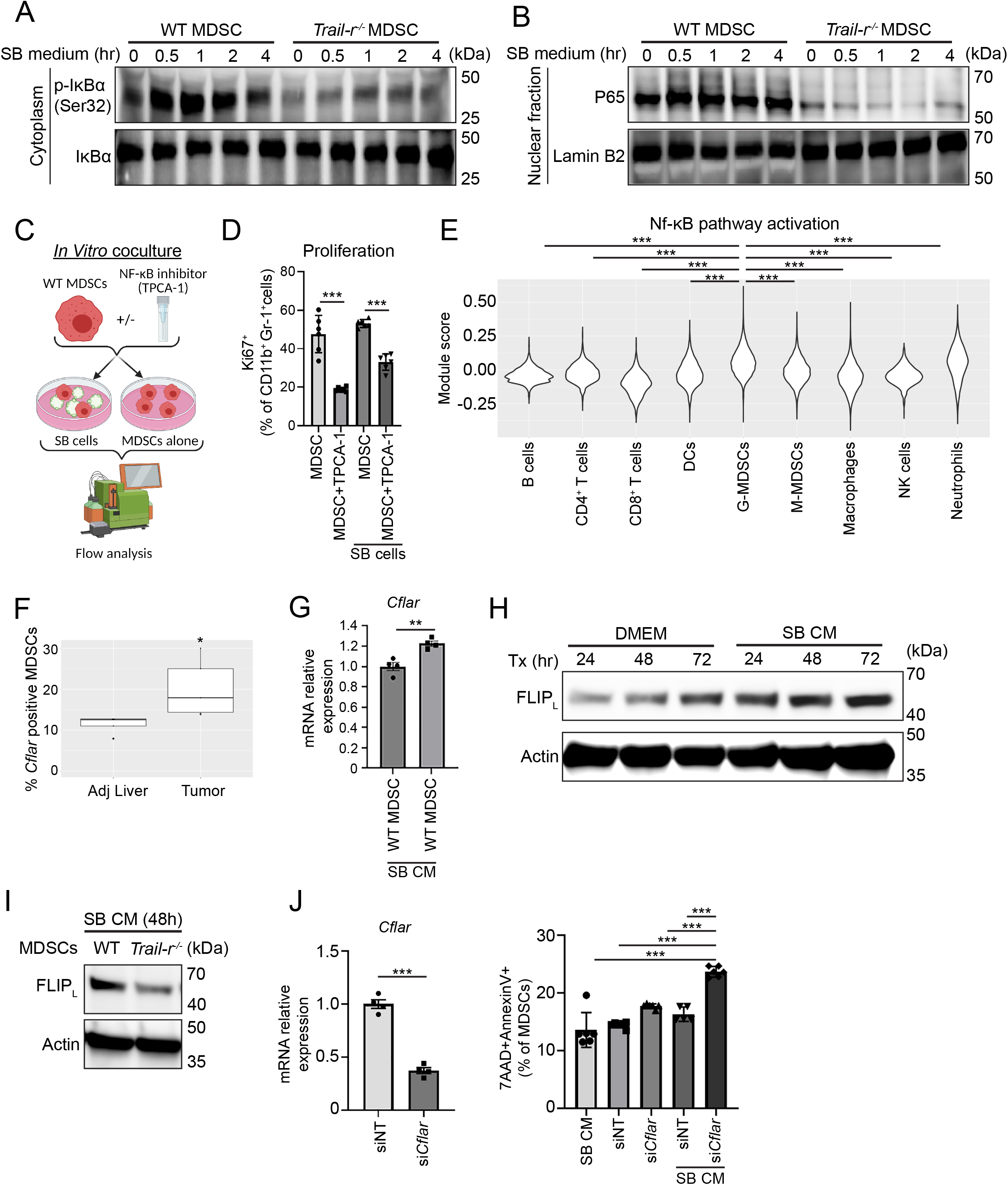
Non-canonical TRAIL-R signaling promotes MDSC survival in an NF-κB dependant manner. (**A**) Immunoblot analysis for IκBα phosphorylation (p-IκBα) in MDSCs isolated from WT and *Trail-r^-/-^*mice and incubated with SB cell conditioned medium. Total IκBα was used as loading control. **(B)** Immunoblot analysis for p65 subunit of NF-κB pathway activation in nuclear extracts of MDSCs differentiated from bone marrow of WT and *Trail-r^-/-^*mice and incubated with SB cell conditioned medium. Lamin B2 was used as loading control. (**C**) Schematic of in vitro experiment with MDSCs differentiated from bone marrow of WT mice and incubated with/without TPCA-1 and/or SB cells. Cells were cocultured for 48 hours. **(D)** Percentage of Ki67^+^ MDSCs (CD11b^+^Gr-1^+^) (*n* ≥ 6). (**E**) Seurat module score for NF-κB pathway^33^ was calculated in tumors from three murine CCA models (SB, KPPC, FAC) using scRNA seq data. The difference of NF-kB pathway activation across different cell types was determined by one-way ANOVA test (*P<*2.2×10^−16^). The difference between two cell types was determined by Tukey multiple comparisons test**. (F)** scRNA seq data demonstrating percentage of *Cflar^+^*MDSCs from tumor-adjacent liver and tumors from SB, KPPC, and FAC tumor bearing mice. **(G)** mRNA expression of *Cflar* in murine WT MDSCs incubated with/without SB CM (n=4). **(H)** Immunoblot analysis for FLIP_L_ in WT murine MDSCs incubated with SB conditioned medium (CM) for 24, 48, and 72 hours. Beta-actin was used as a loading control. **(I)** Immunoblot analysis for FLIP_L_ in murine WT MDSCs and *Trail-r^-/-^* MDSCs after 48h of culture with SB CM. Beta-actin was used as a loading control. **(J)** Percentage of Annexin V^+^7AAD^+^ MDSCs (CD11b^+^Gr-1^+^) following transfection with siNT or si*Cflar* and incubation with/without SB CM for 24 h (*n* ≥ 5). Data are represented as mean ± SD. Unpaired Student’s *t* test was used. \**P* < 0.05; \*\**P* < 0.01; \*\*\**P* < 0.001.

### Enhanced cellular FLICE inhibitory protein (cFLIP) expression confers resistance to canonical, proapoptotic TRAIL signaling in MDSCs

Having demonstrated that MDSCs in murine CCA tumors are resistant to TRAIL-mediated apoptosis **(Fig. 4A)**, we confirmed this finding in vitro as addition of recombinant TRAIL did not alter MDSC viability **(Supplementary Fig. S5D).** Resistance to TRAIL-induced apoptosis can be mediated by inhibitors of death receptor signaling such as cFLIP and antiapoptotic members of the BCL-2 family.^1^ cFLIP is a potent inhibitor of TRAIL proapoptotic signaling.^37^ ScRNA seq and CITE-Seq analysis of murine CCA tumors and adjacent liver demonstrated a significant increase in cFLIP^+^ MDSCs (also known as *Cflar*) in murine tumors compared to tumor-adjacent liver **(Fig. 5F).** To ascertain whether cFLIP expression mediates resistance to TRAIL-mediated apoptosis, murine WT MDSCs were incubated with murine CCA cells. cFLIP RNA and protein expression in MDSCs was significantly increased in the presence of SB cells **(Fig. 5G-H)**. Moreover, compared to WT MDSCs, *Trail-r^-/-^* MDSCs had a significant decrease in cFLIP expression but no change in expression of inhibitors of apoptosis or BCL-2 proteins **(Fig. 5I; Supplementary Fig. S5E)**, suggesting that TRAIL signaling is a major driver of cFLIP expression in MDSCs. Finally, we employed siRNA mediated knockdown of *cFlip* in murine MDSCs (si*Cflar* MDSCs). Compared to non-target (siNT) MDSCs, si*Cflar* MDSCs had a significant increase in apoptosis in the presence of conditioned medium from TRAIL^+^ CCA cells, suggesting that *cFlip* knockdown sensitizes MDSCs to TRAIL-mediated apoptosis **(Fig. 5J)**. Collectively, these findings suggest that enhanced cFLIP expression facilitates MDSC resistance to TRAIL-mediated apoptosis.

### Cancer cell restricted deletion of *Trail* significantly reduces tumor burden

Our data suggest that TRAIL^+^ cancer cells can leverage TRAIL signaling to evade the antitumor immune response. To determine whether targeting TRAIL^+^ cancer cells have therapeutic potential, we employed cancer cell restricted deletion by generating SB cells with CRISPR/Cas9-mediated knockdown of *Trail* (SB-*Trail*^-/-^) and *Trail-r*^-/-^ (SB-*Trail-r*^-/-^) **(Fig. 6A)**. Implantation of SB-*Trail^-/-^* into WT C57BL/6J mice resulted in a significant reduction in the tumor burden compared to control SB cell (SB-NT) implantation **(Fig. 6B-C).** SB-NT, SB-*Trail^-/-^*, SB-*Trail-r*^-/-^ tumors had phenotypic features of CCA **(Supplementary Fig. S6A)**. Tumor-bearing mice were followed with cross-sectional imaging using small animal ultrasound. Ultrasound imaging demonstrated that SB-*Trail^-/-^* tumors were significantly smaller than SB-NT tumors (**(Fig. 6D)**. Concordant with this and similar to our findings in the SB tumor bearing *Trail-r^-/-^* mice, there was a significant decrease in G-MDSCs in the SB-*Trail*^-/-^ tumors without a change in M-MDSCs or macrophages **(Fig. 6E-F; Supplementary Fig. S6B-D)**. Moreover, CTLs and reactive CTLs were also significantly increased in SB-*Trail*^-/-^ tumors **(Fig. 6G-H)**. To confirm that the alteration in MDSC abundance is related to TRAIL-mediated augmentation of MDSC proliferation, CD11b^+^Ly6C^+^ MDSCs were differentiated from BM of WT mice and cocultured with SB-NT and SB-*Trail*^-/-^ cells **(Fig. 6I)**. Consistently, we demonstrated that in the presence of cancer cells devoid of *Trail*, MDSC proliferation was significantly suppressed, while MDSC apoptosis was unchanged **(Fig. 6J)**. These results suggest that cancer cell restricted deletion of *Trail* hinders tumor growth as TRAIL^+^ cancer cells cannot trigger a TRAIL-R mediated increase in MDSC abundance. Notably, SB-*Trail-r^-/-^* cell implantation into WT mice resulted in larger tumors compared to control **(Fig. 6B-C)**. Moreover, SB-*Trail-r^-/-^*tumors had a significant decrease in cancer cell apoptosis compared to SB-NT tumors as assessed by TUNEL staining **(Fig. 6K)**. These results indicate that cancer cell restricted deletion of *Trail-r* has a tumor promoting effect as TRAIL^+^ cytotoxic immune cells cannot trigger TRAIL-R mediated cancer cell apoptosis. In aggregate, these data suggest that TRAIL/TRAIL-R has divergent roles in the tumor immune microenvironment (TIME): TRAIL^+^ cytotoxic immune cells can trigger modest cancer cell death, whereas TRAIL^+^ cancer cells can potently leverage noncanonical TRAIL signaling to evade the antitumor immune response.

**Figure 6.**
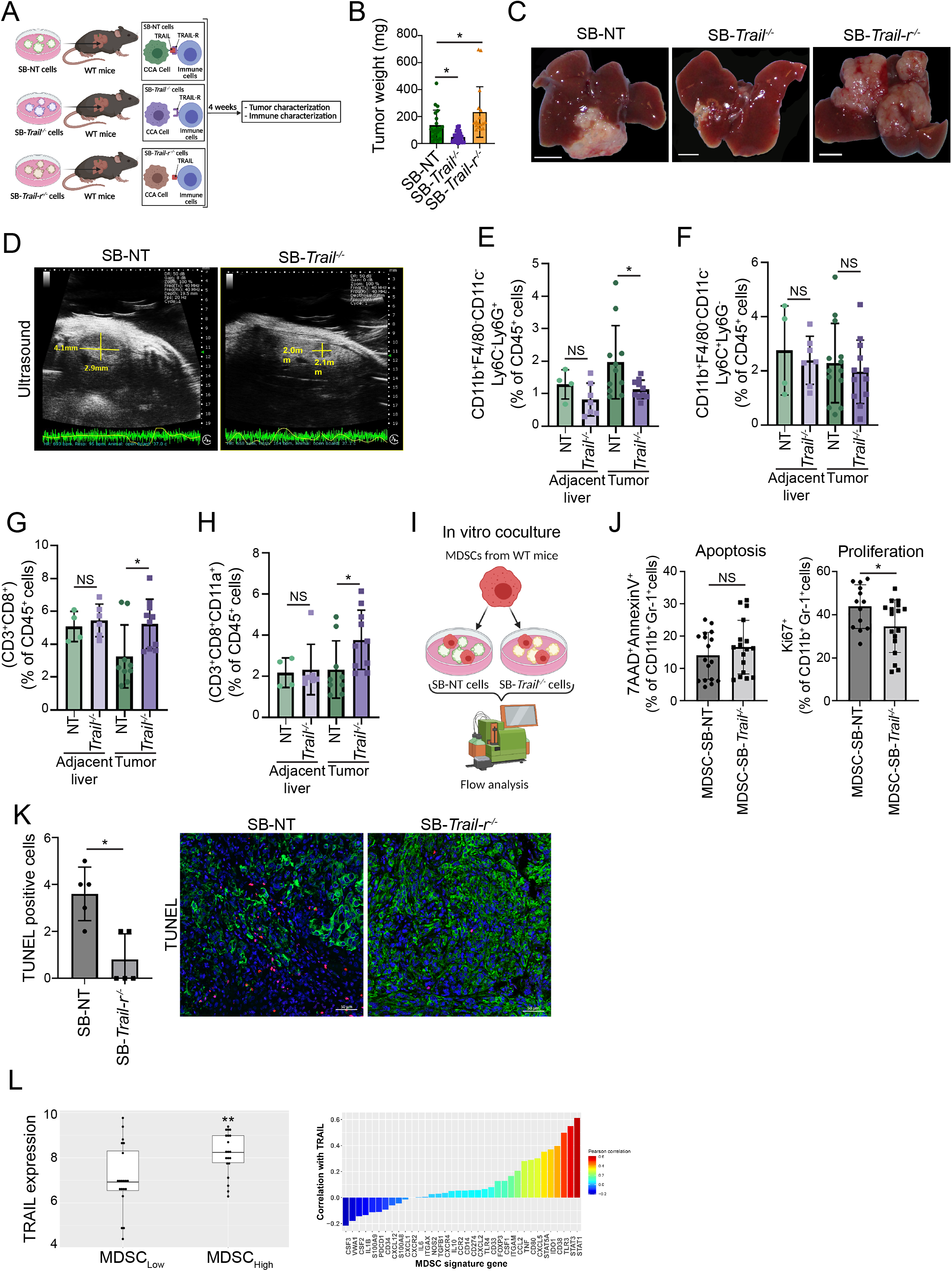
TRAIL/TRAIL-R has pleiotropic effects on the CCA TIME. (**A**) Schematic of WT C57BL/6J mice implanted with SB cells with CRISPR/Cas9-mediated knockout of non-target (SB-NT), *Trail* (SB-*Trail*^-/-^) or TRAIL-R (SB-*Trail-r*^-/-^). (**B**) Average tumor weights in mg of WT mice implanted with SB-NT, SB-*Trail*^-/-^ or SB-*Trail-r*^-/-^ (*n* ≥ 17). (**C**) Representative photographs of livers from **B**. Scale bar: 0.5 cm. (**D**) Ultrasound images depict representative tumor cross-sectional areas on day 25 following implantation of SB-NT or SB-*Trail*^-/-^ cells. Total volumetric area was quantified after 3D reconstruction using Prospect software. (**E**) Percentage of G-MDSCs (CD45^+^F4/80^-^CD11b^+^CD11c^-^Ly6C^-^Ly6G^+^) of CD45^+^ cells in tumor-adjacent livers and tumors from WT mice bearing SB-NT or SB-*Trail*^-/-^ tumors (*n* ≥ 4). (**F**) Percentage of M-MDSCs (CD45^+^ F4/80^-^ CD11b^+^ CD11c^-^ Ly6G^-^ Ly6C^+^) of CD45^+^ cells in tumor-adjacent livers and tumors from WT mice bearing SB-NT or SB-*Trail*^-/-^ tumors (*n* ≥ 4). (**G**) Percentage of CTLs (CD45^+^ CD3^+^ CD8^+^) of CD45^+^ cells in tumor-adjacent livers and tumors from WT mice bearing SB-NT or SB-*Trail*^-/-^ tumors (*n* ≥ 4). (**H**) Percentage of reactive CTLs (CD45^+^ CD3^+^ CD8^+^ CD11a^+^) of CD45^+^ cells in tumor-adjacent livers and tumors from WT mice bearing SB-NT or SB-*Trail*^-/-^ tumors (*n* ≥ 4). (**I**) Schematic depicting in vitro coculture experiment with MDSCs and SB-NT or SB-*Trail*^-/-^ cells. Cells were cocultured for 48h. (**J**) Percentage of Annexin V^+^7AAD^+^ MDSCs (CD11b^+^Gr-1^+^) MDSCs (right panel), and percentage of Ki67^+^ MDSCs (CD11b^+^Gr-1^+^) (left panel) after 48h of coculture with SB-NT or SB-*Trail*^-/-^ cells (*n* ≥ 9). **(K)** Apoptotic cancer cells were quantified by counting TUNEL-positive nuclei in five random microscopic fields using a fluorescent microscope (left panel). Representative photomicrographs depicting apoptotic cancer cells using TUNEL and KRT19 staining (red and green, respectively) in sections of SB-NT or SB-*Trail-r^-/-^* tumors (right panel). Nuclei were counterstained with DAPI. Scale bar: 50 µm. (**L**) The correlation of TRAIL (*TNFSF10*) expression with MDSC signature genes was analyzed using the published Gene Expression Omnibus dataset (GSE76297). Data are represented as mean ± SD. Unpaired Student’s *t* test was used. ns=nonsignificant; \**P* < 0.05; \*\**P* < 0.01.

Finally, to ascertain the human relevance of our findings, we interrogated a gene-expression dataset derived from patients with CCA (GSE76297). Using a previously published MDSC gene signature,^18,38^ we observed that patients with higher MDSC gene expression had significantly higher levels of TRAIL *(TNFSF10)* compared to those with lower MDSC gene expression **(Fig. 6L; Supplementary Figure 6E).** These results are consistent with our preclinical observations that TRAIL augments MDSC abundance.

## DISCUSSION

We have identified a potent immunosuppressive role of TRAIL signaling in the CCA TIME. These data indicate that i) TRAIL/TRAIL-R actively supports tumor growth as *Trail-r* deficiency across multiple immunocompetent models of CCA significantly reduces tumor volumes; ii) TRAIL signaling promotes tumor immunosuppression by increasing MDSC accumulation in murine CCA tumors; iii) noncanonical TRAIL signaling augments MDSC abundance via enhanced proliferation in an NF-κB dependent manner; iv) MDSCs are resistant to TRAIL-mediated apoptosis due to enhanced cFLIP expression; v) cancer cell restricted deletion of *Trail* significantly reduces tumor burden via decreased MDSC abundance.

Proapoptotic TRAIL signaling as a cause of cancer cell death is a well-established mechanism, whereby TRAIL^+^ cytotoxic immune cells, such as NK cells, bind to TRAIL-R on cancer cells with consequent cancer cell apoptosis. However, TRAIL-R agonists have had very limited anticancer activity in human clinical trials.^1^ The existing literature indicates that the effects of TRAIL signaling in the TIME are pleiotropic and context dependent. Whole body *Trail-r* deficiency in several murine models of tumorigenesis (e.g., breast cancer, thymoma) renders mice more susceptible to primary tumor growth and metastasis.^10,11^ In contrast, TRAIL induced cytokine secretion from TRAIL apoptosis resistant cancer cells promotes a tumor supportive TIME in lung adenocarcinoma.^5^ This TRAIL-triggered secretome promotes MDSCs and TAMs, indicating a tumor promoting immune modulatory role of TRAIL signaling.^5^ Similarly, our data has demonstrated a significant reduction in CCA tumor burden of *Trail-r* deficient mice. However, the TRAIL paradigm described in lung adenocarcinoma^5^ is intact in our model, as the CCA cells implanted into mouse livers express both *Trail* and *Trail-r* while the immune cells lack *Trail-r*. This suggests an alternate mechanism for TRAIL induced increase in MDSCs and a concomitant decrease in CTLs in the CCA TIME.

The TRAIL/TRAIL-R system has been implicated in T cell homeostasis.^30^ TRAIL^+^ astrocytes maintain CNS homeostasis by inducing T cell apoptosis.^39^ In experimental models of autoimmune diseases, engagement of TRAIL-R decreases CD8^+^ T cell proliferation.^40^ Thus, we postulated that TRAIL/TRAIL-R may directly impact CTL apoptosis or proliferation. However, in vitro functional studies did not demonstrate a TRAIL-mediated alteration in T cell apoptosis or proliferation. Indeed, conditional deletion of *Trail-r* in CD8^+^ T cells did not impact tumor volumes. Moreover, myeloid cell restricted deletion of *Trail-r* resulted in an increase in CTL abundance in murine CCA tumors, likely due to an indirect effect via modulation of immunosuppressive MDSC abundance by TRAIL/TRAIL-R. MDSCs are immature myeloid cells that mediate tumor immune escape with consequent tumor progression.^18,19,41^ An abundance of MDSCs in the TIME is associated with resistance to ICI and chemotherapy.^41-43^ Although MDSCs are pivotal in tumor immunosuppression, therapeutic strategies targeting MDSCs are limited.^41^ Moreover, the available myeloid modulators have had limited success in clinical trials.^41^

In the current study, we have identified a cell autonomous effect of noncanonical TRAIL signaling in MDSCs. Noncanonical TRAIL signaling cascades can predominate in certain contexts, promoting proinflammatory gene transcription programs and activation of NF-κB,^13,44^ but the role of noncanonical TRAIL signaling in tumor immunology is not clear.^1,30,45^ In cells that are resistant to TRAIL-induced apoptosis, NF-κB activation via noncanonical TRAIL signaling can induce proliferation.^44^ For instance, TRAIL signaling can induce proliferation of TRAIL apoptosis resistant tumor cells in an NF-κB dependent manner.^6,44^ Our study has found that this paradigm also exists in MDSCs as WT MDSCs had NF-κB activation that was abrogated in *Trail-r^-/-^*MDSCs, and NF-κB inhibition attenuated TRAIL-mediated increase in MDSC proliferation. Moreover, single cell transcriptomics demonstrated that MDSCs across multiple murine CCA models had significant enrichment of an NF-κB activation signature.

It is well-established that enhanced cFLIP expression promotes prosurvival, noncanonical TRAIL signaling and confers resistance to canonical, proapoptotic TRAIL signaling in cancer cells.^46-49^ cFLIP also protects M-MDSCs from chemotherapy induced death and promotes immunosuppressive activity in M-MDSCs.^50^ Moreover, pharmacological targeting of cFLIP can sensitize cancer cells to proapoptotic TRAIL signaling.^48,51^ In congruence with these prior observations, our findings suggest that cFLIP also plays an essential role in mediating MDSC resistance to TRAIL-mediated apoptosis. WT MDSCs had enhanced cFLIP expression compared to *Trail-r^-/-^*MDSCs. Moreover, cFLIP knockdown sensitized MDSCs to TRAIL-mediated apoptosis, suggesting that cFLIP protects MDSCs from cell death. Thus, our studies suggest that switching prosurvival/proliferation TRAIL signaling to canonical, proapoptotic TRAIL signaling will promote MDSC apoptosis, which in turn has therapeutic implications for tumor suppression. The combination of immunotherapeutics (e.g., ICI) and TRAIL-targeting therapies has been assessed in preclinical models of cancer.^52-55^ However, these combinatorial approaches have employed TRAIL agonism, rather than selective TRAIL antagonism on cancer cells. We have now demonstrated that TRAIL/TRAIL-R system has a dual role in the tumor microenvironment by facilitating cancer cell apoptosis (canonical) while promoting an immunosuppressive tumor microenvironment by directly fostering MDSC abundance and function (noncanonical). Cancer cell restricted deletion of *Trail* significantly reduced MDSC accumulation and enhanced T cell cytotoxicity with a consequent reduction in tumor burden. Conversely, cancer cell restricted deletion of *Trail-r* increased murine tumor growth by promoting MDSC abundance and function. These observations highlight that systemic inhibition of TRAIL/TRAIL-R may lead to unwanted attenuation of apoptosis of TRAIL-R^+^ CCA cells by TRAIL^+^ immune cells. In summary, our findings support the role of selective therapeutic targeting of TRAIL^+^ cancer cells in an effort to block TRAIL/TRAIL-R mediated tumor immunosuppression.

## METHODS

### Mice

All animal experiments were performed in accordance with a protocol approved by the Mayo Clinic Institutional Animal Care and Use Committee (IACUC). Eight-week-old male C57BL/6J mice, and LysM^cre^ C57BL/6J mice were purchased from Jackson Laboratories. LysM^cre^ mice are myeloid-cre expression mice with high recombination rates in myeloid cells including mature macrophage and MDSCs.^32^ *Trail-r^f/f^*, *Trail^f/f^* and *Trail-r^-/-^* were bred at Mayo Clinic. *LysM^cre^-Trail-r^f/f^* mice were bred at Mayo Clinic.

### Syngeneic, orthotopic murine models of CCA

We have established multiple, syngeneic immunocompetent models of CCA that recapitulate human CCA and have distinct genetic drivers.^25-27^ The SB murine CCA cell line and the resultant syngeneic model^26^ are derived from a genetic model employing biliary instillation of the oncogenes *Akt* and yes-associated protein (*YAP^S127A^*).^25^ This Akt/YAP driven model has substantial overlap with human iCCA.^25^ Mutations inducing activation of the oncogene *Kras* and inactivation of the tumor suppressor gene *p53* are common genetic alteration in CCA.^56,57^ *Kras^G12D^p53^L/L^*murine CCA tumors (KPPC) are more aggressive than SB tumors and have a distinct phenotype representing a sarcomatoid CCA subtype, a rare and aggressive subtype of human iCCA driven by *Kras^G12D^p53^L/L^*.^58^ In contrast, implantation of CCA cells (FAC) derived from a genetic model driven by loss of *FBXW7* and permissive Akt^59^, results in a syngeneic model with slower tumor growth than the SB model (manuscript in preparation). Murine CCA cells (SB, KPPC, FAC)^25,27,59^ were maintained in culture medium as previously described.^26^ Mice were anesthetized and 20 µL of standard media containing 0.75 x 10^6^ SB, 0.75 x 10^6^ KPPC, or 1.0x10^6^ FAC cells were injected orthotopically into the lateral aspect of the medial lobe as previously described.^18,26^ Mice were sacrificed two, four, and five weeks, respectively, after implantation of KPPC, SB, and FAC cells. Tumor, adjacent liver, and spleen were harvested following sacrifice.

### Generation of SB-*Trail^-/-^* and SB-*Trail-r^-/-^* cells

CRISPR/Cas9-mediated knockout of mouse *Trail* and *Trail-r* in SB cells was conducted using the dual single guide RNA vector system pSpCas9(bb)-2A-GFP (PX458). The sgRNA sequences sgRNA1-*Trail-r*, sgRNA2-*Trail-r*, forward-*Trail-r*, reverse-*Trail-r*, sgRNA1-*Trail*, sgRNA2-*Trail*, forward-*Trail* and reverse-*Trail* were generated in PZH vector amplified and extracted using In-Fusion HD Cloning kit (Clontech). Plasmids were sent for Sanger sequencing with reverse and forward primer to control any mutation point. SB cells were then transfected using lipofectamine 3000 transfection kit (Invitrogen). SB clone knockout for *Trail* or *Trail-r* were selected by PCR.

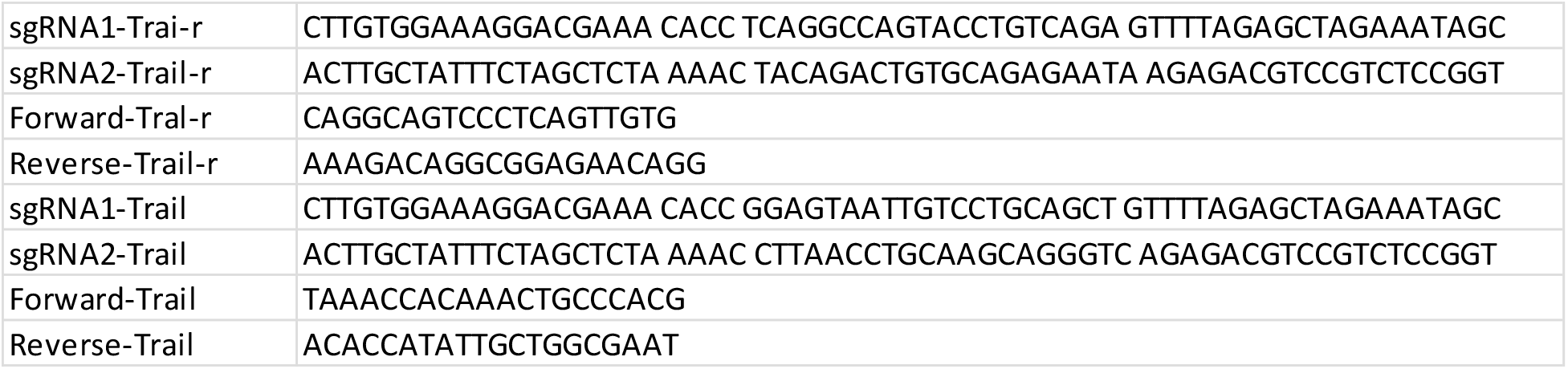

### Isolation of liver, spleen, or tumor-infiltrating immune cells and flow analysis

Upon excision, tumor and liver were dissociated with gentleMACS^™^ Octo Dissociator (Miltenyi) according to the manufacturer’s protocol and as previously described.^18^ CD45^+^ cells were isolated by CD45 (TIL) mouse microbeads (Miltenyi). Cells were stained and data was acquired on a Miltenyi MACSQuant® Analyzer 10 optical bench flow cytometer as previously described.^18^ All antibodies were used following the manufacturer recommendation. Fluorescence Minus One controls were used for each independent experiment to establish gates. For intracellular staining of granzyme B, CD206, IL-10, Ki67 and Arginase1, cells were stained using the intracellular staining kit (Miltenyi). Apoptosis staining was performed using 7AAD-annexinV-APC kit (BioLegend®). Data analysis was conducted using FlowJo™ (TreeStar). Forward scatter (FSC) and side scatter (SSC) were used to exclude cell debris and doublets. Fixable Viability Stain 510 (BD Horizon™) was used to exclude dead cells.

### Isolation of bone marrow MDSCs (BM-MDSC) and in vitro culture

BM-MDSCs were isolated from the legs of WT, *Trail-r^-/-^*, *Trail^f/f^*, and *LysM^cre^-Trail-r^f/f^* mice as previously described.^33^ Bone marrow was extracted and filtered through a 70 μm cell strainer. Single cell suspension was cultured in MDSC activation medium (RPMI 1640, 10% low endotoxin FBS, interleukin [IL]-6 40 ng/ml, granulocyte colony-stimulating factor 40 ng/ml, granulocyte-macrophage colony-stimulating factor 40 ng/ml) for 5 days to induce MDSC differentiation. BM-MDSC were plated in 24-well plate at 1x10^5^ cells and co-cultured with SB cells at a 1:4 ratio or treated with TPCA-1 (GW683965) 10 µM for 24h. BM-MDSC were stained with Fixable Viability Stain 510 (BD Horizon™), CD11b-PE-Cy5, Gr-1-PE, PD-L1-BV421, IL-10-FITC, Arginase1-PE-Cy7, Ki67-AF700, AnnexinV-APC, 7-AAD and analyzed by flow cytometry.

### Small animal ultrasound imaging

Mice were anesthetized using 1.5–3% isoflurane via a nose cone. Abdominal hair was removed using clippers followed by depilatory cream. Transabdominal ultrasound in B mode was recorded using the Prospect T1 high-frequency ultrasound system (S-Sharp Corporation) with a 40 MHz transducer. 2D and 3D morphometric parameters were analyzed offline.

### Sample preparation for scRNA seq and CITE-Seq

Adjacent liver and tumor samples were collected from SB, KPPC, and FAC tumor bearing C57BL/6J mice. The samples were minced and dissociated into single cells using the Miltenyi dissociation kit (mouse tumor dissociation kit [Miltenyi 130-096-730]) for tumor samples or mouse liver dissociation kit [Miltenyi 130-105-807] for adjacent uninvolved liver samples). Cell digestion of both tumor and adjacent normal samples was strained through a 70 μm cell strainer, and blood cells were eliminated using a red blood cell lysis buffer. Single cells were stained with viability dye (BD Horizon 564406) and PE/Cyanine7 antimouse CD45 Antibody (BioLegend® 103113). CD45 positive viable cells were sorted using the BD FACSMelody™ Cell Sorter. Cells were stained with TotalSeq™-C Mouse Universal Cocktail (BioLegend® V1.0, 199903). Finally, the stained single cells suspension was processed and sequenced at the Genome Analysis Core.

### Gene module analysis

To evaluate the activity of the NF-κB pathway, the score for NF-κB was calculated with AddModuleScore function from Seurat package using the BioCarta NF-κB pathway gene set.^36^

### Statistical analysis

Experimental statistical analyses were performed using GraphPad Prism software. Student two-tailed *t* test, log-rank (Mantel–Cox) test, and one-way ANOVA (with Tukey multiple comparison test) were used throughout as indicated in the text. Data were considered statistically significant at *P* < 0.05.

## Abbreviations

CCA: cholangiocarcinoma
CITE-Seq: cellular indexing of transcriptomes and epitopes by sequencing
cFLIP: cellular FLICE inhibitory protein
CTL: cytotoxic T lymphocyte
G-MDSCs: granulocytic-MDSCs
ICI: immune checkpoint inhibition
MDSCs: myeloid-derived suppressor cells
M-MDSCs: monocytic-MDSCs
NK: natural killer cells
PD-1: programmed death-1
PD-L1: programmed death-ligand 1
scRNA seq: single cell RNA sequencing
TAMs: tumor-associated macrophages
TIME: tumor immune microenvironment
TRAIL: Tumor necrosis factor related apoptosis inducing ligand

## Acknowledgments

We thank Ms. Jenn Rud for excellent administrative support. KPPC cells were kindly gifted by Nabeel Bardeesy, PhD (Harvard Medical School, Massachusetts General Hospital, Broad Institute). This publication was made possible through support by core facilities at the Mayo Clinic including Medical Genome Facility Gene Analysis Core, Pathology Research Core, Optical Microscopy Core, and the Antibody Hybridoma Core.

