## Supplementary Information for "Noncanonical TRAIL Signaling Promotes Myeloid-Derived Suppressor Cell Abundance and Tumor Progression in Cholangiocarcinoma"

### Fig. S1

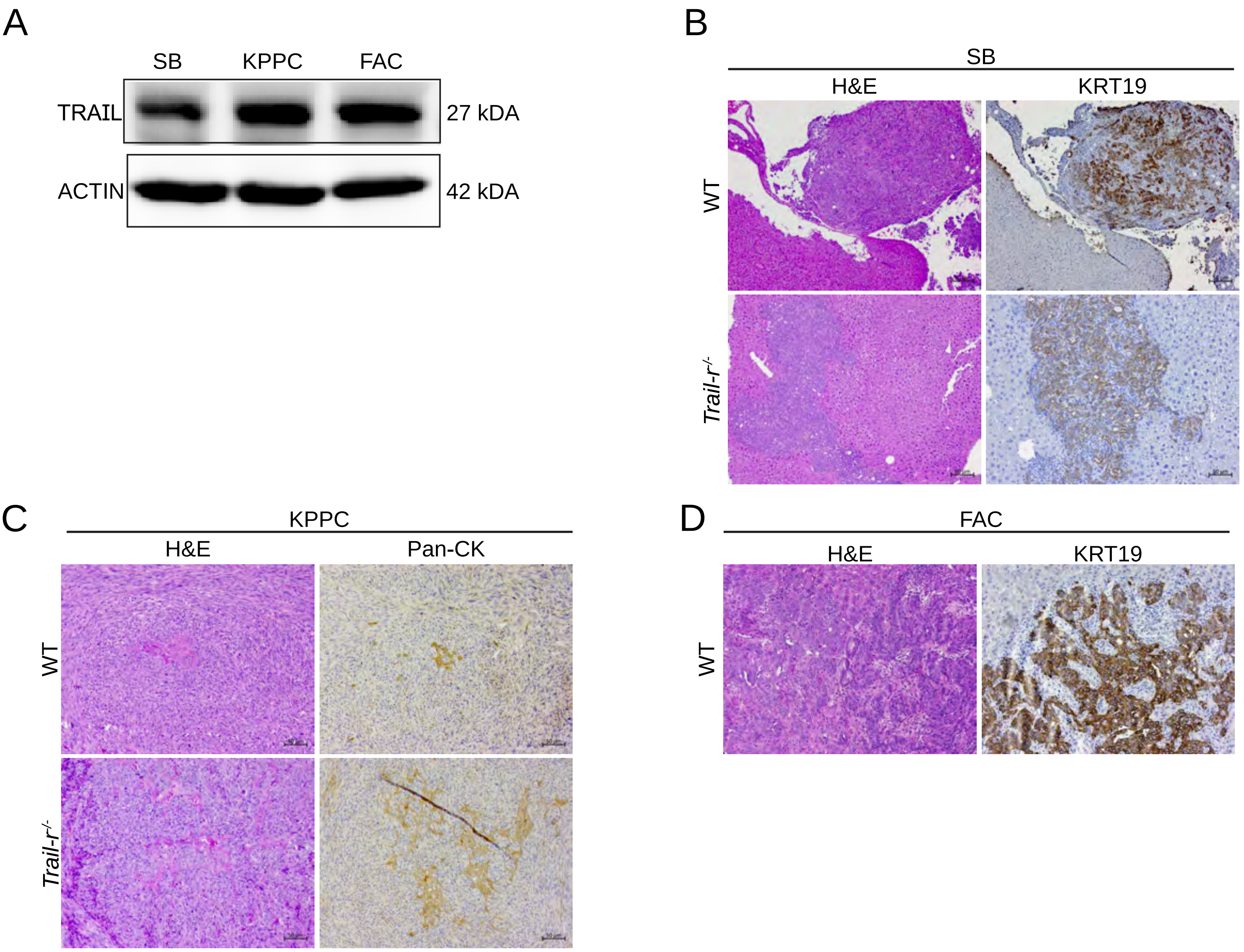
A
SB
KPPC
FAC
TRAIL
27 kDA
ACTIN
42 kDA
B
SB
H&E
KRT19
WT
Trail-r-/-
C
KPPC
H&E
Pan-CK
WT
Trail-r-/-
D
FAC
H&E
KRT19
WT

Fig. S2

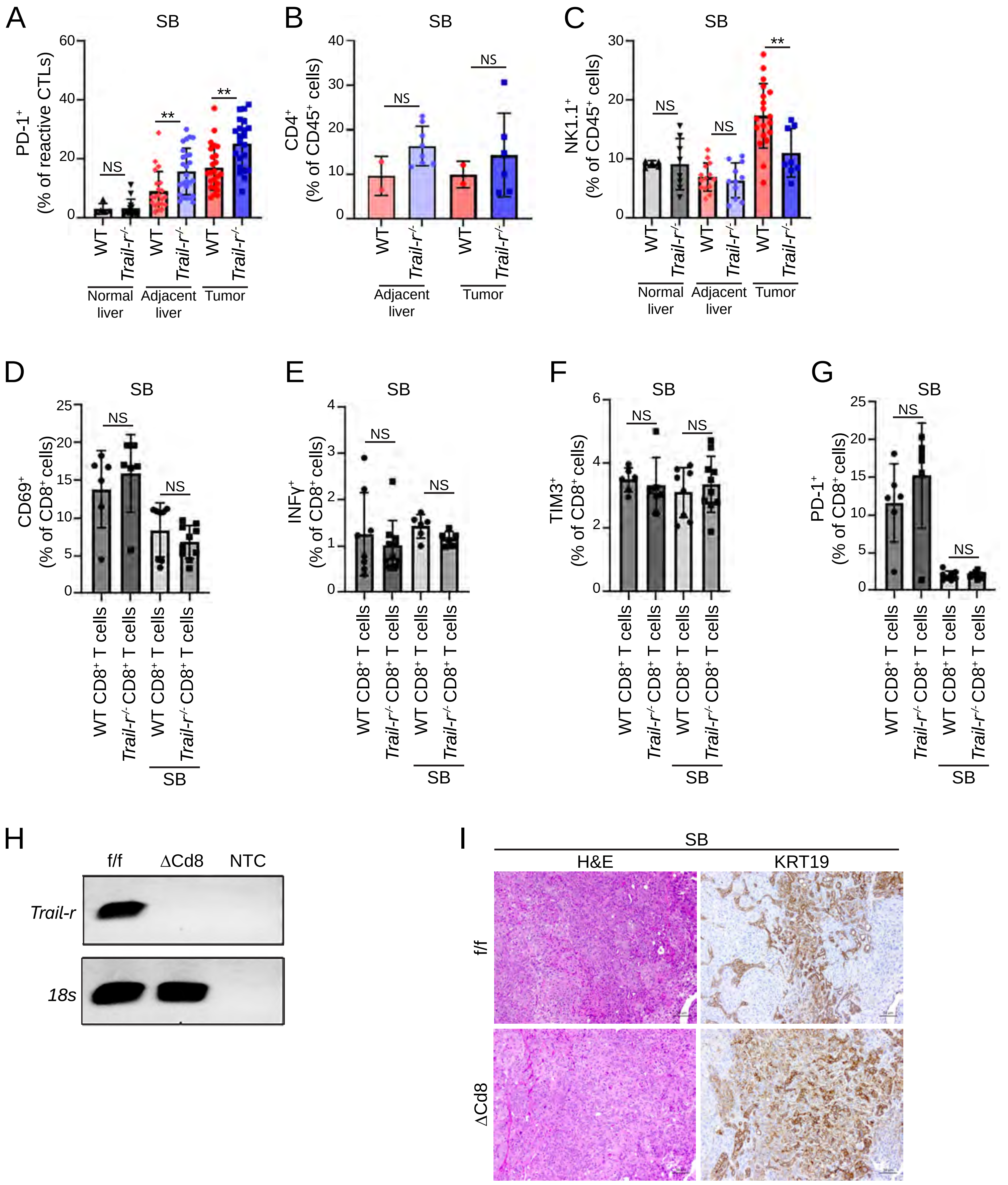
A
SB
PD-1+ (% of reactive CTLs)
60
40
20
0
NS
**
**
WT
Trail-r-/-
WT
Trail-r-/-
WT
Trail-r-/-
Normal liver
Adjacent liver
Tumor
B
SB
CD4+ (% of CD45+ cells)
40
30
20
10
0
NS
NS
WT
Trail-r-/-
WT
Trail-r-/-
Adjacent liver
Tumor
C
SB
NK1.1+ (% of CD45+ cells)
30
20
10
0
NS
NS
**
WT
Trail-r-/-
WT
Trail-r-/-
WT
Trail-r-/-
Normal liver
Adjacent liver
Tumor
D
SB
CD69+ (% of CD8+ cells)
25
20
15
10
5
0
NS
NS
WT CD8+ T cells
Trail-r-/- CD8+ T cells
WT CD8+ T cells
Trail-r-/- CD8+ T cells
SB
E
SB
INFγ+ (% of CD8+ cells)
4
3
2
1
0
NS
NS
WT CD8+ T cells
Trail-r-/- CD8+ T cells
WT CD8+ T cells
Trail-r-/- CD8+ T cells
SB
F
SB
TIM3+ (% of CD8+ cells)
6
4
2
0
NS
NS
WT CD8+ T cells
Trail-r-/- CD8+ T cells
WT CD8+ T cells
Trail-r-/- CD8+ T cells
SB
G
SB
PD-1+ (% of CD8+ cells)
25
20
15
10
5
0
NS
NS
WT CD8+ T cells
Trail-r-/- CD8+ T cells
WT CD8+ T cells
Trail-r-/- CD8+ T cells
SB
H
f/f
ΔCd8
NTC
Trail-r
18s
I
SB
H&E
KRT19
f/f
ΔCd8

Fig. S3

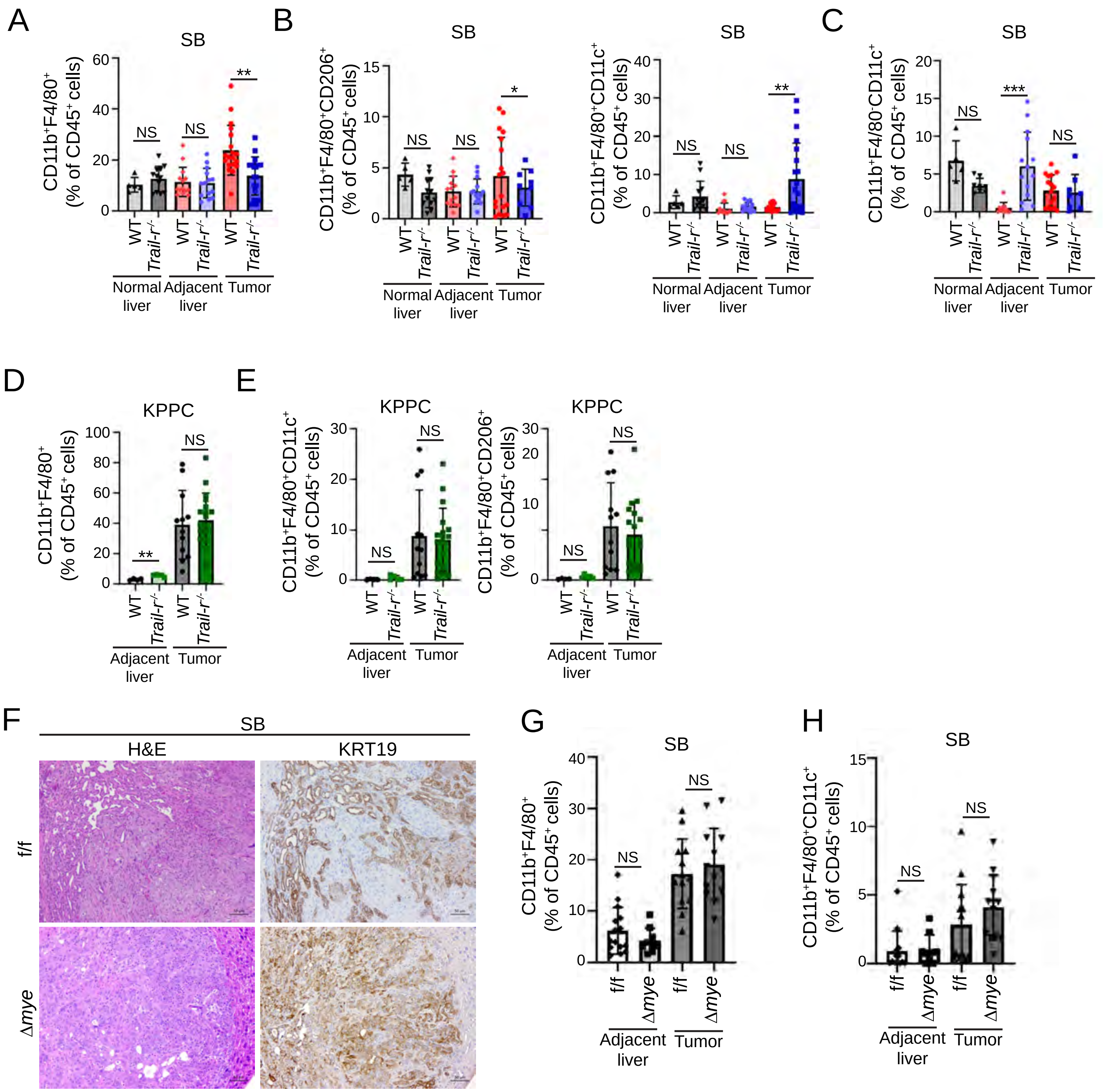
A
SB
CD11b+F4/80+ (% of CD45+ cells)
60
40
20
0
NS
NS
**
WT
Trail-r-/-
WT
Trail-r-/-
WT
Trail-r-/-
Normal liver
Adjacent liver
Tumor
B
SB
CD11b+F4/80+CD206+ (% of CD45+ cells)
15
10
5
0
NS
NS
*
SB
CD11b+F4/80+CD11c+ (% of CD45+ cells)
40
30
20
10
0
NS
NS
**
C
SB
CD11b+F4/80-CD11c+ (% of CD45+ cells)
20
15
10
5
0
NS
***
NS
D
KPPC
CD11b+F4/80+ (% of CD45+ cells)
100
80
60
40
20
0
**
NS
Adjacent liver
Tumor
E
KPPC
CD11b+F4/80+CD11c+ (% of CD45+ cells)
30
20
10
0
NS
NS
KPPC
CD11b+F4/80+CD206+ (% of CD45+ cells)
30
20
10
0
NS
NS
F
SB
H&E
KRT19
f/f
Δmye
G
SB
CD11b+F4/80+ (% of CD45+ cells)
40
30
20
10
0
NS
NS
f/f
Δmye
f/f
Δmye
Adjacent liver
Tumor
H
SB
CD11b+F4/80+CD11c+ (% of CD45+ cells)
15
10
5
0
NS
NS
f/f
Δmye
f/f
Δmye
Adjacent liver
Tumor

Fig. S4

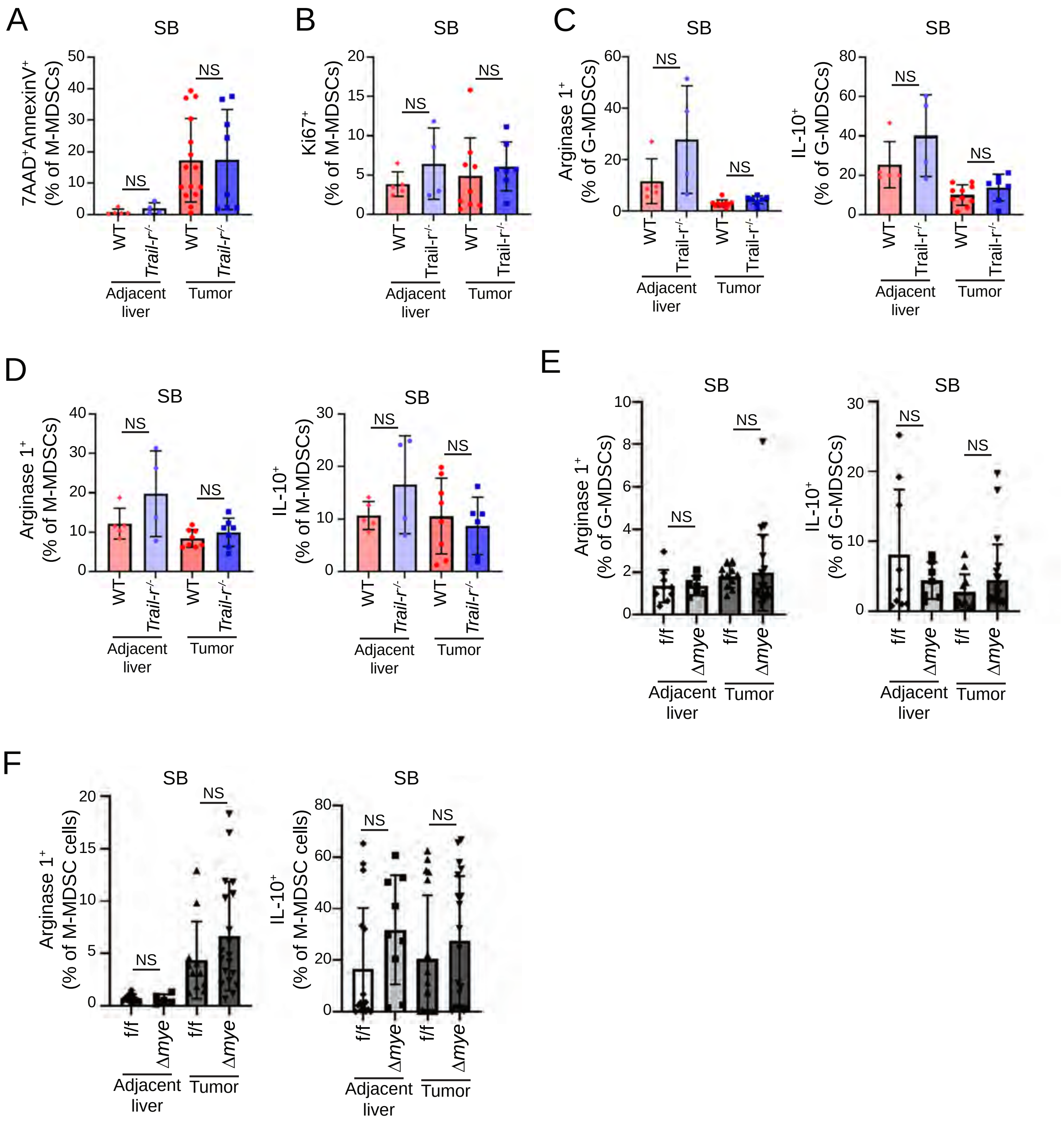
A
SB
7AAD⁺AnnexinV⁺ (% of M-MDSCs)
NS
NS
WT
Trail-r-/-
WT
Trail-r-/-
Adjacent liver
Tumor
B
SB
Ki67⁺ (% of M-MDSCs)
NS
NS
Adjacent liver
Tumor
C
SB
Arginase 1⁺ (% of G-MDSCs)
NS
NS
SB
IL-10⁺ (% of G-MDSCs)
NS
NS
D
SB
Arginase 1⁺ (% of M-MDSCs)
NS
NS
SB
IL-10⁺ (% of M-MDSCs)
NS
NS
E
SB
Arginase 1⁺ (% of G-MDSCs)
NS
NS
f/f
Δmye
f/f
Δmye
Adjacent liver
Tumor
SB
IL-10⁺ (% of G-MDSCs)
NS
NS
F
SB
Arginase 1⁺ (% of M-MDSC cells)
NS
NS
SB
IL-10⁺ (% of M-MDSC cells)
NS
NS

### Figure S5

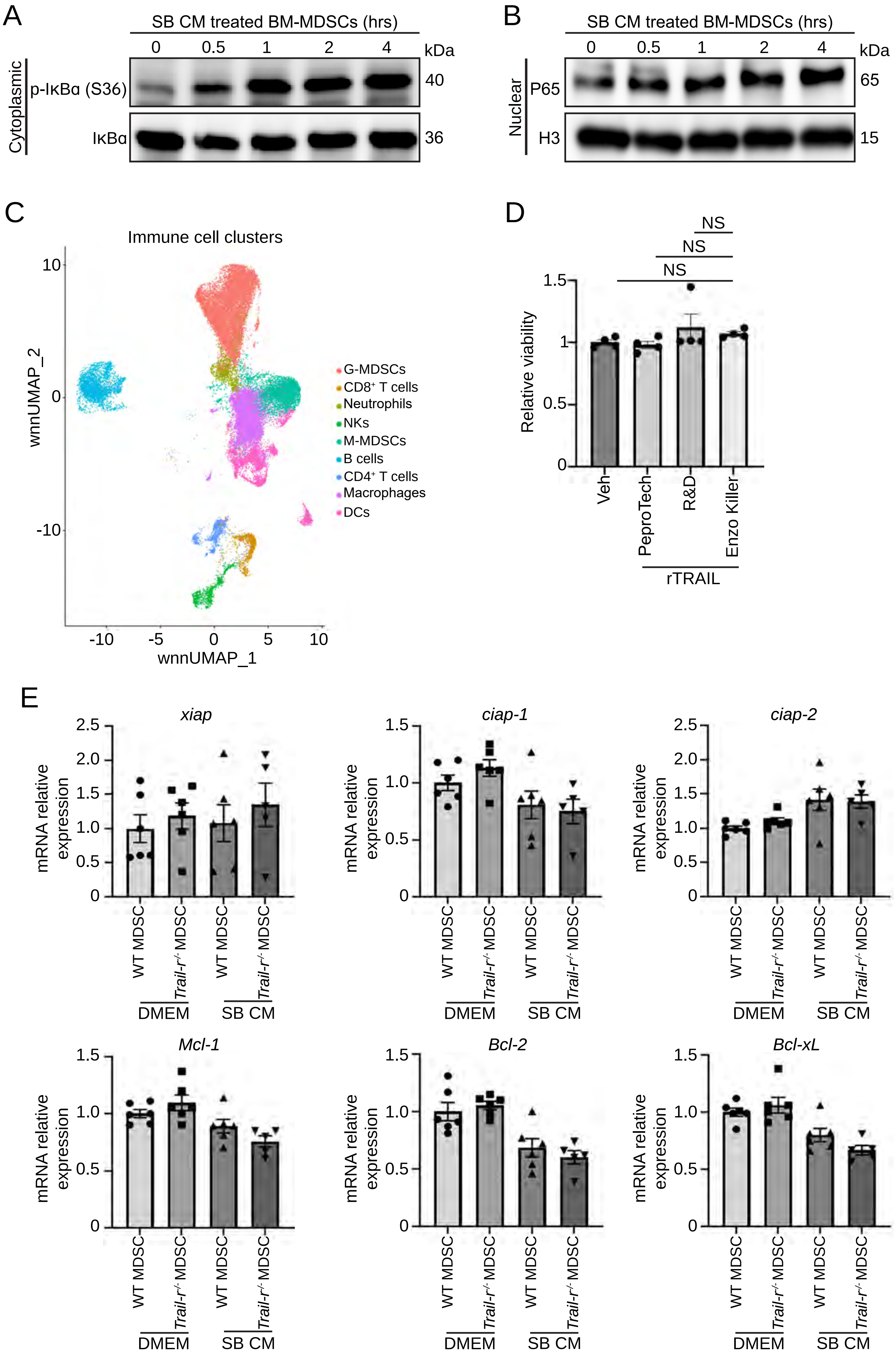
A
SB CM treated BM-MDSCs (hrs)
0
0.5
1
2
4
kDa
Cytoplasmic
p-IκBα (S36)
40
IκBα
36
B
SB CM treated BM-MDSCs (hrs)
0
0.5
1
2
4
kDa
Nuclear
P65
65
H3
15
C
Immune cell clusters
wnnUMAP_2
wnnUMAP_1
G-MDSCs
CD8+ T cells
Neutrophils
NKs
M-MDSCs
B cells
CD4+ T cells
Macrophages
DCs
D
NS
NS
NS
Relative viability
Veh
PeproTech
R&D
Enzo Killer
rTRAIL
E
xiap
ciap-1
ciap-2
Mcl-1
Bcl-2
Bcl-xL
mRNA relative expression
WT MDSC
Trail-r-/- MDSC
WT MDSC
Trail-r-/- MDSC
DMEM
SB CM

### Figure S6

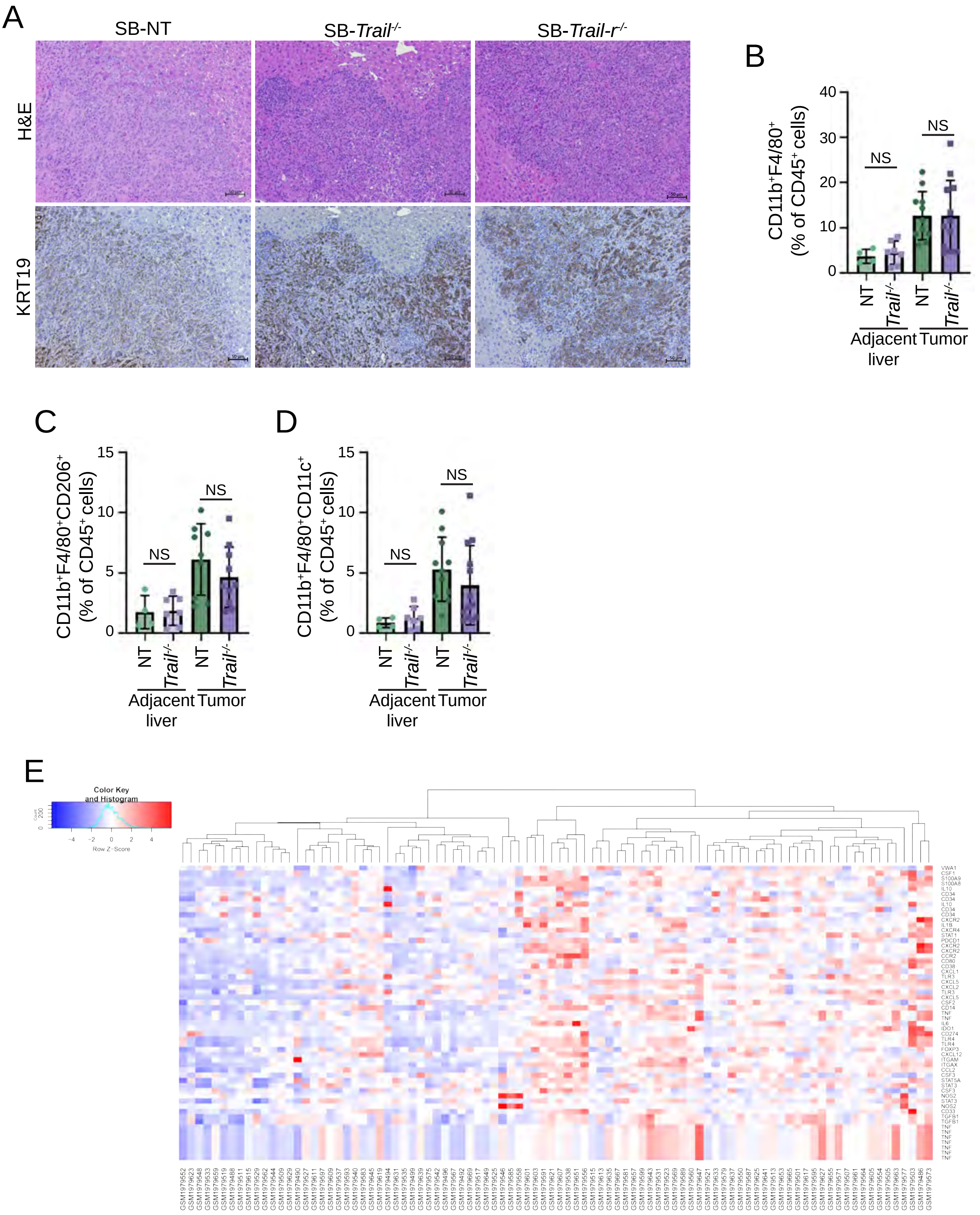
A
SB-NT
SB-Trail-/-
SB-Trail-r-/-
H&E
KRT19
B
CD11b+F4/80+ (% of CD45+ cells)
NS
NS
NT
Trail-/-
NT
Trail-/-
Adjacent liver
Tumor
C
CD11b+F4/80+CD206+ (% of CD45+ cells)
NS
NS
Adjacent liver
Tumor
D
CD11b+F4/80+CD11c+ (% of CD45+ cells)
NS
NS
Adjacent liver
Tumor
E
Color Key and Histogram
Count
Row Z-Score

#### SUPPLEMENTARY FIGURE LEGENDS

**Supplementary Figure S1.** (**A**) Immunoblot analysis of TRAIL expression in SB, KPPC and FAC mouse CCA cell lysates. Beta-actin was used as a loading control. (**B**) Representative photomicrographs of hematoxylin and eosin-stained (left panel) and KRT19 immunohistochemistry (IHC) (right panel) of SB tumor sections from WT and *Trail-r*$^{-/-}$ mice. Scale bar: 50 µm. (**C**) Representative photomicrographs of hematoxylin and eosin-stained (left panel) and pan-keratin IHC (right panel) of KPPC tumor sections from WT and *Trail-r*$^{-/-}$ mice. Scale bar: 50 µm. (**D**) Representative photomicrographs of hematoxylin and eosin-stained (left panel) and KRT19 IHC (right panel) of FAC tumor sections from WT mice. Scale bar: 50 µm.

**Supplementary Figure S2.** (**A–C**) Tumor growth of 28 days after orthotopic implantation of $0.75\times10^6$ SB cells in WT or *Trail-r*$^{-/-}$ mouse livers. (**A**) Percentage of PD-1$^+$ reactive CTLs (CD45$^+$ CD3$^+$ CD8$^+$CD11a$^+$)) in WT mouse livers from mice without tumors, tumor-adjacent livers and SB tumors from WT and *Trail-r*$^{-/-}$ mice ($n \geq 5$). (**B**) Percentage of CD4$^+$ T cells (CD45$^+$CD3$^+$CD4$^+$) of CD45$^+$ cells in tumor-adjacent livers and SB tumors from WT and *Trail-r*$^{-/-}$ mice ($n \geq 5$). (**C**) Percentage of natural killer (NK) cells (CD45$^+$CD3$^-$NK1.1$^+$) of CD45$^+$ cells in tumor-adjacent livers and SB tumors from WT and *Trail-r*$^{-/-}$ mice ($n \geq 5$). (**D**) Percentage of CD69$^+$ CTLs (CD45$^+$CD3$^+$CD8$^+$) following 36h of coculture with SB cells ($n \geq 5$). (**E**) Percentage of INFγ$^+$ CTLs (CD45$^+$CD3$^+$CD8$^+$) following 36h of coculture with SB cells ($n \geq 5$). (**F**) Percentage of TIM3$^+$ CTLs (CD45$^+$CD3$^+$CD8$^+$) following 36h of coculture with SB cells ($n \geq 5$). (**G**) Percentage of PD-1$^+$ CTLs (CD45$^+$CD3$^+$CD8$^+$) following 36h of coculture with SB cells ($n \geq 5$). (**H**) Expression of *Trail-r* in CD8$^+$ T cells isolated from *Trail-r*$^{f/f}$ (f/f) and *CD8*$^{cre}$*Trail-r*$^{f/f}$ (ΔCd8) mice. 18s rRNA was used as a normalization control. Water was used as a no template control (NTC). (**I**) Representative photomicrographs of hematoxylin and eosin stained and KRT19 IHC of SB tumor section from f/f and ΔCd8 mice. Scale bar: 50 µm. Data are represented as mean ± SD. Unpaired Student's *t* test was used. ns=nonsignificant; $^{**}P < 0.01$.

**Supplementary Figure S3.** (**A–C**) Immune profiling following tumor growth of 28 days after orthotopic implantation of $7.5\times10^5$ SB cells in WT or *Trail-r*$^{-/-}$ mouse livers. (**A**) Percentage of macrophages (CD45$^+$F4/80$^+$CD11b$^+$) of CD45$^+$ cells (right panel) in WT mouse livers from mice without tumors, tumor-adjacent livers and SB tumors from WT and *Trail-r*$^{-/-}$ mice ($n \geq 5$). (**B**) Percentage of TAMs (CD45$^+$F4/80$^+$CD11b$^+$CD206$^+$) (left panel) and M1-like macrophages (CD45$^+$F4/80$^+$CD11b$^+$CD11c$^+$) (right panel) of CD45$^+$ cells in WT mouse livers from mice without tumors, tumor-adjacent livers and SB tumors from WT and *Trail-r*$^{-/-}$ mice ($n \geq 5$). (**C**) Percentage of dendritic cells ($CD45^+F4/80^-CD11b^+CD11c^+$) of $CD45^+$ cells in WT mouse livers from mice without tumors, tumor-adjacent livers and SB tumors from WT and *Trail-r*$^{-/-}$ mice ($n \geq 5$). (**D–E**) Immune profiling following tumor growth of 14 days after orthotopic implantation $7.5 \times 10^5$ KPPC cells in WT or *Trail-r*$^{-/-}$ mouse livers. (**D**) Percentage of macrophages ($CD45^+F4/80^+CD11b^+$) of $CD45^+$ cells in tumor-adjacent livers and KPPC tumors from WT and *Trail-r*$^{-/-}$ mice ($n \geq 4$). (**E**) Percentage of M1-like macrophages ($CD45^+F4/80^+CD11b^+$ $CD11c^+$) (left panel) and TAMs ($CD45^+F4/80^+CD11b^+CD206^+$) (right panel) of $CD45^+$ cells in tumor-adjacent livers and KPPC tumors from WT and *Trail-r*$^{-/-}$ mice ($n \geq 4$). (**F–H**) Tumor growth of 28 days after orthotopic implantation of $0.75 \times 10^6$ SB cells in f/f and Δmye mouse livers. (**F**) Representative photomicrographs of hematoxylin and eosin-stained (left panel) and KRT19 IHC (right panel) of SB tumor sections from f/f and Δmye mice. Scale bar: 50 µm. (**G**) Percentage of macrophages ($CD45^+F4/80^+CD11b^+$) $CD45^+$ cells in tumor-adjacent livers and SB tumors from f/fand Δmye mice ($n \geq 6$). (**H**) Percentage of M1-like macrophages ($CD45^+F4/80^+CD11b^+CD11c^+$) of $CD45^+$ cells in tumor-adjacent livers and SB tumors from f/f and Δmye mice ($n \geq 6$). Data are represented as mean ± SD. Unpaired Student's *t* test was used. ns=nonsignificant; *$P < 0.05$; **$P < 0.01$; ***$P < 0.001$.

**Supplementary Figure S4.** (**A**) Percentage of Annexin $V^+7AAD^+$ M-MDSCs ($CD45^+F4/80^-$ $CD11b^+CD11c^-Ly6C^+Ly6G^-$) of $CD45^+$ cells in tumor-adjacent livers and SB tumors from WT and *Trail-r*$^{-/-}$ mice. (**B**) Percentage of $Ki67^+$ M-MDSCs ($CD45^+F4/80^-CD11b^+CD11c^-$ $Ly6C^+Ly6G^-$) of $CD45^+$ cells in tumor-adjacent livers and SB tumors from WT and *Trail-r*$^{-/-}$ mice ($n \geq 4$). (**C**) Percentage of Arginase $1^+$ G-MDSCs (left panel) and IL-$10^+$ G-MDSCs (right panel) in tumor-adjacent livers and SB tumors from WT and *Trail-r*$^{-/-}$ mice ($n \geq 4$). (**D**) Percentage of Arginase $1^+$ M-MDSCs (left panel) and IL-$10^+$ M-MDSCs (right panel) in tumor-adjacent livers and SB tumors from WT and *Trail-r*$^{-/-}$ mice ($n \geq 4$). (**E**) Percentage of Arginase $1^+$ G-MDSCs (left panel) and IL-$10^+$ G-MDSCs (right panel) in tumor-adjacent livers and SB tumors from f/f and Δmye mice ($n \geq 6$). (**F**) Percentage of Arginase $1^+$ M-MDSCs (left panel) and IL-$10^+$ M-MDSCs (right panel) in tumor-adjacent livers and SB tumors from f/f and Δmye mice ($n \geq 6$). Data are represented as mean ± SD. Unpaired Student's *t* test was used. ns=nonsignificant.

**Supplementary Figure S5. (A)** Immunoblot analysis for IκBα phosphorylation in BM-MDSCs incubated with SB cell CM. Total IκBα was used as a loading control. **(B)** Immunoblot analysis for p65 subunit of NF-κB pathway activation in nuclear extracts of BM-MDSCs incubated with SB cell conditioned medium. Histone H3 was used as loading control. (**C**) mRNA expression of *xiap*, *ciap-1*, *ciap-2*, Mcl-1, *Bcl-2*, and *Bcl-xL* in WT and *Trail-r*$^{-/-}$ MDSCs incubated with control medium (DMEM) or SB conditioned medium (SB CM). (**D**) UMAP plot of 42031 immune cells from SB, KPPC, and FAC tumors. (**E**) Viability assay in BM-MDSCs treated with vehicle or different types of recombinant mouse TRAIL (200 ng/ml) for 48 hours (n=4). Data are represented as mean ± SD. Unpaired Student's *t* test was used. ns=nonsignificant.

**Supplementary Figure S6.** (**A**) Representative photomicrographs of hematoxylin and eosin-stained (left panel) and KRT19 IHC (right panel) of SB tumor sections from WT mice implanted with SB-NT, SB-*Trail*$^{-/-}$ or SB-*Trail-r*$^{-/-}$ cells. (**B**) Percentage of macrophages (CD45$^{+}$F4/80$^{+}$CD11b$^{+}$) of CD45$^{+}$ cells in tumor-adjacent livers and SB-NT or SB-*Trail*$^{-/-}$ tumors ($n \geq 4$). (**C**) Percentage of TAMs (CD45$^{+}$ F4/80$^{+}$ CD11b$^{+}$ CD206$^{+}$) of CD45$^{+}$ cells in tumor-adjacent livers and SB-NT or SB-*Trail*$^{-/-}$ tumors ($n \geq 4$). (**D**) Percentage of M1-like macrophages (CD45$^{+}$ F4/80$^{+}$ CD11b$^{+}$ CD11c$^{+}$) of CD45$^{+}$ cells in tumor-adjacent livers and SB-NT or SB-*Trail*$^{-/-}$ tumors ($n \geq 4$). (**E**) Heatmap depicting normalized gene expression data for MDSC signature genes for 92 CCA tumor samples from Gene Expression Omnibus dataset (GSE76297). Data are represented as mean ± SD. Unpaired Student's *t* test was used. ns=nonsignificant.

#### SUPPLEMENTARY METHODS

*Cell culture and treatment*

All cells were maintained in culture at 37ºC and 5% $CO_2$ in Dulbecco's Modified Eagle Medium (DMEM; Thermo Fisher Scientific) supplemented with 10% (v/v) fetal bovine serum (FBS; Thermo Fisher Scientific), 0.2% (v/v) primocin (Thermo Fisher Scientific), and 1% penicillin (100 IU/mL; Thermo Fisher Scientific), and streptomycin (100 µg/mL; Thermo Fisher Scientific). SB cells were previously generated by us.[1] FAC cells were also generated by us (manuscript in preparation). KPPC cells were kindly gifted by Nabeel Bardeesy, PhD (Harvard Medical School, Massachusetts General Hospital, Broad Institute).[2,3] The following siRNAs at 10 nM final concentration were used: N-TARGETplus Non-targeting Control Pool (horizon, # D-001810-10-20), ON-TARGETplus Mouse Cflar siRNA SMARTPool (horizon, # L-041091-00-0005). Lipofectamine™ RNAiMAX Transfection Reagent (ThermoFisher, # 13778150) was used for siRNA transfection. Recombinant mouse TRAILs were purchased from PeproTech (#315-19), R&D Systems (#1121-TL-010) and Enzo Life Sciences (#ALX-201-073-C020). TPCA-1 (Selleckchem, #S2824) at 10 µM final concentration was used. CellTiter-Glo® 2.0 Cell Viability Assay (Promega, # G9242) was used for viability assay per manufacturers' protocol.

*Isolation of T cells*

$CD8^+$ T cells were isolated from a single cell suspension of WT, *Trail*$^{f/f}$, *CD8*$^{cre}$-*Trail-r*$^{f/f}$ or *Trail-r*$^{-/-}$ mouse spleens as previously described.[18] Briefly, single cell suspension was filtered through 70 µm mesh, centrifuged at 400 g for 10 minutes and re-suspended at $1x10^8$ cell/ml. $CD8^+$ T cells were isolated using the EasySep™ Mouse CD8+ T Cell Isolation Kit (#19853, STEMCELL) per the manufacturer's protocol.

*Flow cytometry antibodies*

The following antibodies were used for flow cytometry staining: F4/80-PacificBlue (BM8, BioLegend®), CD11b-PE-Cy5 (M1/70, eBioscience™), CD11b-PE-Cy7 (M1/70, BioLegend®)

CD206-PE-Cy7 (C068C2, BioLegend®), CD11c-APC (REA754, Miltenyi), Gr-1-PE (RB6-8C5, BioLegend®), Ly6G-PE (REA526, Miltenyi), Ly6C-APC-Vio®770 (REA796, Miltenyi), Arginase1-PE-Cy7 (A1exF5, eBioscience™), IL-10-FITC (JES5-16E3, BioLegend®), Ki67-AF700 (16A8, BioLegend®), PD-L1-BV421 (10F.9G2, BioLegend®), CD3-APC-Cy7 (17A2, BioLegend®), CD8-BV421 (53-6.7, BD Horizon™), CD11a-PE-Vio®770 (REA880, Miltenyi), PD-1-PerCP-Vio®700 (REA802, Miltenyi), granzyme B-PE (QA16A02, BioLegend®), NK1.1-APC (PK136, Miltenyi), CD69-PE-Cy7 (H1.2F3, BioLegend®), TIM-3-FITC (REA602, Miltenyi), INFγ-PE (XMG1.2, BioLegend®) and 7AAD-annexinV-APC kit (640930, BioLegend®).

*Immunoblot analysis*

For the SB conditioned medium studies, BM-MDSC were plated in 10 cm dishes, and on the following day were incubated with equal volume of SB cell conditioned medium for 0, 0.5, 1, 2 or 4h. BM-MDSC cytoplasmic and nuclear protein lysates were collected using the NE-PER™ Nuclear and Cytoplasmic Extraction Reagents (ThermoFisher, #78833) per the manufacturer's protocol. Proteins were resolved by SDS-PAGE and transferred to nitrocellulose membranes. The following primary antibodies were used for immunoblotting analysis: polyclonal TRAIL (Abcam, #ab231265, or ab42121-100), IκBα (Cell Signaling Technology, #4814s), Phospho-IκBα (Ser32) (Cell Signaling Technology, #2859s), Phospho-IκBα (Ser36) (Abcam, #ab133462), NF-κB p65 (Santa Cruz, #SC-8008), Lamin B2 (Cell Signaling Technology, # #12255), Histone H3 (Cell signaling Technology, #4499s), cFLIP (Cell Signaling Technology, #5634s). Membranes were blotted with primary antibody overnight at 4°C at a dilution of 1:1000. Goat Anti-Mouse or Rabbit IgG (H + L)-HRP Conjugate secondary antibodies were used at a dilution of 1:2000. Proteins were visualized with Clarity and Clarity Max ECL Western Blotting Substrates (Bio-Rad).

*Immunohistochemistry in mouse tumor and adjacent liver specimens*

Tumor and adjacent liver tissue from euthanized mice was fixed in 4% formalin for 48h, embedded in paraffin, and sectioned into 3–5 µm slices. Formalin-fixed, paraffin-embedded mouse tumor and adjacent liver sections were deparaffinized, hydrated and incubated with primary antibody overnight at 4°C. Sections were stained with antibody for keratin 19 (KRT19) (Cell Signaling, 12434, 1:500) or Pan-Keratin (Cell signaling, C11, 1:500). Primary antibody was detected with HRP-conjugated secondary antibody and diaminobenzidine (Dako, Carpentaria, CA, USA). Liver tissue sections were counterstained with hematoxylin or methyl green.

*Reverse transcription, polymerase chain reaction (PCR) and Real-time PCR*

BM-MDSCs at day 5 were isolated as described above and treated as indicated in figure legend. RNA was extracted using TRIzol™ Reagent (ThermoFisher, #15596018), and cDNA was generated using M-MuLV Reverse Transcriptase (NEB, #M0253L). The following primers targeting the cDNA sequence were used for genotyping: *Trail-r* forward: 5`-CCCAGCCCATAATCGTCCAG-`3`; *Trail-r* reverse: 5`-TCCAGAGAATGGTTGGAATGGC-3`. *18S* forward: 5`-CGCTTCCTTACCTGGTGGAT -3`; *18S* reverse: 5`-GAGCGACCAAAGGAACCATA-3`. The following quantitative PCR primers were used: *Cflar* forward: 5`-GCTCCAGAATGGGCGAAGTAA-3`; *Cflar* reverse: 5`-ACGGATGTGCGGAGGTAAAAA-3`. *Xiap* forward: 5`- CGAGCTGGGTTTCTTTATACCG-3`; *Xiap* reverse: 5`-GCAATTTGGGGATATTCTCCTGT-3`. *Ciap1* forward: 5`-TGTGGCCTGATGTTGGATAAC-3`; *Ciap1* reverse: 5`-GGTGACGAATGTGCAAATCTACT-3`. *Mcl-1* forward: 5`-AAAGGCGGCTGCATAAGTC-3`; *Mcl-1* reverse: 5`-TGGCGGTATAGGTCGTCCTC-3`. *Bcl2* forward: 5`-ATGCTGCAAAACGTGACTCC-3`; *Bcl-2* reverse: 5`-AAGCTGGTACAGAAGCCATTG-3`.

*Bcl-xl* forward: 5`-GACAAGGAGATGCAGGTATTGG-3`; *Bcl-xl* reverse: 5`-TCCCGTAGAGATCCACAAAAGT-3`. *Gapdh* forward: 5`-AGGTCGGTGTGAACGGATTTG-3`; *Gapdh* reverse: 5`-TGTAGACCATGTAGTTGAGGTCA-3`.

*TUNEL Assay in Mouse Tumor Specimens*

The fluorescent TUNEL assay was performed using the *In Situ* Cell Death Detection Kit (Roche Diagnostics) on formalin-fixed paraffin-embedded tissue sections. Briefly, tissues were deparaffinized and rehydrated, with antigen retrieval achieved via hot citrate buffer. The tissues were then stained with the TUNEL reaction mixture according to the manufacturer's instructions. In addition, the tissues were incubated with primary antibody for KRT19 (Cell Signaling) at a concentration of 1:100 overnight at 4°C. Nuclei were counterstained with DAPI. Apoptotic tumor cells were quantified by counting TUNEL-positive, KRT19 positive nuclei in five random microscopic fields per slide (at 10x magnification) using a LSM780 confocal microscope (Zeiss). Mann-Whitney test was used to determine significance.

*Analysis of public scRNA seq datasets*

The raw reads of scRNAseq data of tumors from 13 treatment-naive intrahepatic cholangiocarcinoma (iCCA) patients[4] were downloaded from the Genome Sequence Archive (HRA000863). Transcriptomic reads were aligned to the GRCh38 reference genome and quantified using Cellranger (v3.0). Gene count matrix from scRNA seq data of tumors from 4 patients with biliary tract cancer undergoing surgical resection[5] was downloaded from GEO accession GSE210066. Gene count matrix from scRNA seq data of tumors from 3 patients with iCCA was downloaded from GEO accession GSE189903.[6] Gene count matrix from scRNA seq data of tumors from 4 patients with iCCA was downloaded from GEO accession GSE138709.[7]

Gene count matrix from scRNA seq data of tumors from 10 iCCA patients[8] was downloaded from GEO accession GSE125449. The gene counts data was further processed using Seurat v4.0. Cells were selected for downstead analysis if they met the following quality control criteria: i) gene counts ≤ 3 mean absolute deviation (MAD) above median and > 200; ii) < 5% ratio of mitochondrial genes. Gene counts data was normalized using SCT transformation and integrated using harmony v0.1.1 to eliminate batch effects. PCA was used for dimension reduction and the shared nearest neighbor (SSN) algorithm was employed to construct the shared nearest neighbor graph using the first 50 principal components. A total of 41 clusters were identified with unsupervised clustering using Louvain community detection at a resolution of 0.8. The following cell markers were used to identify the major cell types: CCL21, VWF, ACKR1, SPARCL1 and PECAM1 for endothelial cell; KRT19, KRT7, EPCAM, KRT18, SOX9 and ANXA4 for epithelial cells; COL1A1, COL1A2, COL3A1, ACTA2 and IGFBP7 for fibroblasts; CD3D, CD3E, CD3G and TRAC for T cells; NKG7, GNLY, NCAM1 and KLRD1 for NK cells; CD79A, IGHM, IGHG3 and IGHA2 for B cells; LYZ, MARCO, CD68 and FCGR3A for myeloid cells.

*Analysis of public RNA sequencing data*

The normalized counts data was downloaded from GEO datasets (GSE76297) and filtered for tumor samples in CCA patients. A total of 92 CCA tumor samples were identified and used for downstream analysis. A heatmap was used to depict the expression of MDSC signature genes in each sample.[9] Based on the order of the samples in heatmap, the 20 samples with the highest expression of MDSC signature genes were defined as the $MDSC_{high}$ group. The 20 samples with the lowest expression of MDSC signature genes were defined as the $MDSC_{low}$ group. The expression of TRAIL (*TNFSF10)* was compared between the $MDSC_{high}$ group and $MDSC_{low}$ group.

*scRNA seq and CITE-seq data processing*

For samples with CITE-seq data, the raw reads from both RNA and protein were aligned to the GRCh38 reference genome and quantified using Cellranger (v3.0). The gene counts matrix was further processed using Seurat v4.0. Doublet cells were identified and removed using DoubletFinder v2. Dead cells were filtered out based on a criterion that required the ratio of mitochondrial genes to be lower than 5%. To ensure data normalization, SCT transformation and CLR normalization were applied to RNA data and protein data, respectively. To eliminate batch effects, the data from multiple samples were integrated with RPCA for RNA and protein, respectively. After data integration, the weighted nearest neighbor (WNN) algorithm was employed to construct the shared nearest neighbor graph using both RNA and protein data. The cells were grouped to 35 clusters by unsupervised clustering using Louvain community detection at resolution of 2. The cell types were identified by examining the expression of the following cell type markers: Cd19, Ighd and Cd27 for B cell, Cd3e, Cd8a, Cd8b1 and Cd4 for T cells, Itga2, Cd3e and Klrb1c for NK cells, Ly6g, Ly6c1 and Arg1 for MDSCs; Adgre1 and Cd68 for macrophages; Itgax, H2, Bst2, Siglech, and Cd83 for dendritic cells; Itgam, Csf1r, Ly6, Ly6c1, Ly6g and Adgre1 for neutrophils, Cd200r3 and Fcer1a for basophils.

For the samples with only scRNA data, the raw reads were aligned to the GRCh38 reference genome and quantified using Cellranger (v3.0). The gene counts matrix was further processed using Seurat v4.0. Doublet cells were identified and removed using DoubletFinder v2. Dead cells were filtered out based on a criterion that required the ratio of mitochondrial genes to be lower than 5%. SCT transformation was applied to RNA data to ensure data normalization. Finally, the cell types were classified by referencing to the CITE-seq data using MapQuery function from Seurat package.
